# Convergent stochastic assembly governs reef biofilm microbiomes across ecologically distinct benthic substrates

**DOI:** 10.64898/2026.09.02.748686

**Authors:** Jordan A. Sims, Jennifer L. Salerno

## Abstract

Understanding the processes that shape microbial biodiversity and community structure is a key objective of the field of microbial ecology. The processes driving assembly of benthic biofilm bacteria on functionally important reef substrates, such as crustose coralline algae (CCA) and calcium carbonate, are not well understood, despite their critical contributions to the maintenance of biodiversity and ecosystem function on reefs. To characterize the patterns of community assembly and biogeography on these substrates, climax biofilm bacterial communities from 11 reef sites were collected, and full 16S small subunit rRNA genes were sequenced. Though CCA- and carbonate-associated communities demonstrated different diversity, composition, and correlations with environmental conditions, communities on both substrates were assembled according to similar processes. Stochastic processes dominated assembly on both substrates, primarily drift with moderate influence from dispersal limitation and selection. Sub-communities of habitat generalists and specialists, as well as rare and abundant taxa, experienced disparate patterns of assembly that remained consistent between substrates, highlighting the importance of individual taxa traits in shaping community assembly. These results provide insight into the factors shaping benthic biofilm bacterial assembly and biogeography in a tropical reef ecosystem and contribute to understanding of reef resilience in the face of environmental change.

## 1. INTRODUCTION

Large-scale ecosystem functioning on coral reefs is driven in part by many activities occurring on a <1 millimeter scale, deemed micro-scale processes (Mullen et al. 2016). Benthic microbial biofilms play a role in many of these micro-scale processes, including nutrient fluxes to and from the seafloor and the settlement and recruitment of corals and numerous other sessile invertebrates (Hadfield & Paul 2001, Fabricius 2005). These functions allow the spatial distribution of biofilm taxa to influence the composition and structure, and thereby function, of the large-scale adult communities on reefs in appreciable ways. Despite the critical ecological role they play, benthic biofilms on reefs are poorly characterized in the literature, and little is known about the biogeography of reef biofilms or the processes that shape these microbial communities.

Microbial communities had historically been theorized to be shaped only by environmental factors due to their potential to disperse widely (Baas-Becking 1934). However, developments in sequencing technology have improved the ability to observe and characterize microbial biogeography and have subsequently advanced ecological theory to indicate microbial community structure may also arise through interactive competing processes (Chave 2004), borrowing from both niche and neutral ecological theory. In niche theory, different microbial taxa are assumed to possess unique traits (Leibold 1995); in combination with environmental variability supporting an infinite number of potential niches, extremely high microbial diversity arises (Leibold & McPeek 2006). In this model, microbial distribution and diversity are controlled by deterministic processes. Neutral theory, conversely, treats taxa as ecologically equivalent with uniform fitness, so diversity is determined by stochastic processes (Hubbell 2001, Chave 2004).

The theoretical framework established by Vellend (2010) theorizes that community assembly arises through the influence of four ecological processes: selection, drift, dispersal, and diversification. These processes are parallel to the evolutionary concepts of selection, genetic drift, gene flow, and mutation, respectively, which shape populations of individual taxa (Hanson et al. 2012). In this study, we use these terms as defined by Nemergut et al. (2013), as follows.

Selection refers to changes in microbial community structure caused by deterministic fitness differences between taxa, a definitionally deterministic process. Selection in the face of low variability in environmental conditions or biotic interactions will drive communities to be more similar and is termed “homogeneous selection”, while selection under highly heterogeneous biotic or abiotic drivers over space or time will drive communities to be more different and is called “heterogeneous” or “variable” selection (Zhou & Ning 2017). Drift refers to random changes in the relative abundances of different microbial taxa within a community through time and is inherently a stochastic process. Dispersal refers to the movement and successful establishment of organisms across space, with high dispersal rates that increase similarly among distinct communities considered to be “homogenizing dispersal” and low dispersal rates that drive differences in disparate communities considered to be “dispersal limitation” (Zhou & Ning 2017). Diversification refers to the generation of new genetic variation through processes such as speciation and horizontal gene transfer. Both dispersal and diversification are considered largely stochastic in microbial communities, but under certain conditions, these processes may be shaped by deterministic components (Zhou & Ning 2017).

This theoretical framework is increasingly being applied to study microbial communities from diverse ecosystems, such as soil (Dumbrell et al. 2010), groundwater (Graham et al. 2017), and montane lakes (Liao et al. 2016). In most cases, both stochastic and deterministic processes are found to contribute to community assembly and biogeography, and a statistical framework has since been iteratively developed to parse and quantify the relative roles of these different assembly processes (Stegen et al. 2013, 2015). This framework assesses (1) phylogenetic turnover to assign the role of selection and (2) taxonomic turnover to assign the roles of dispersal and drift (Zhou & Ning 2017), and it was later extended to be performed on phylogenetically similar bins, rather than whole communities, which further increased its accuracy, precision, sensitivity, and specificity (Ning et al. 2020).

Despite the regular use of this statistical framework in terrestrial environments, studies using this method in marine environments are rare and mostly limited to seawater and sediments (Liu et al. 2019). In general, planktonic marine microbes are structured primarily by stochastic processes, with deterministic processes playing a secondary role, but the exact interactions are highly dependent on the type of microbial taxa investigated (i.e., bacteria, archaea, or microeukaryotes) and environmental conditions such as depth and nutrient concentration (Dai et al. 2017, Chen et al. 2017, Wu et al. 2018, Mo et al. 2018, Wang et al. 2019, Zhang et al. 2021, Ma et al. 2022, Zhao et al. 2024). These observations are possibly due to homogenization of microbial communities and environmental conditions by ocean currents (Liu et al. 2019). Studies from marine sediments tell a different story, often demonstrating strong influence from deterministic rather than stochastic processes in bacterial and archaeal community assembly and highlighting pH as an important environmental driver of selection (Song et al. 2021, Ma et al. 2022, Gong et al. 2022, Cheng et al. 2023, Xu et al. 2024). Even fewer studies have used this framework in coral reef ecosystems. One study analyzed seawater from the reefs of the Xisha Islands and showed strong influence of stochastic structuring, similar to other studies of planktonic communities (Liu et al. 2022). Two others used the framework to investigate assembly of the coral microbiome and found that it is strongly influenced by selection and dispersal limitation, indicating that the coral host plays an active role in structuring its microbiome (Zhang et al. 2021, Zhao et al. 2024).

In addition to analyses of the whole microbial community, subcommunity-level analyses can be performed on taxa with similar traits. Subcommunities of habitat generalists and specialists or rare and abundant taxa are regularly found to follow different patterns of assembly and biogeography than the microbial community as a whole. Habitat generalists are often studied in the context of environmental change because they are more resistant than specialists and may support the persistence of community function in the face of environmental disturbance (Chen et al. 2021). In marine sediments, community assembly of generalists has been shown to be dominated by stochastic processes, with generalists demonstrating high adaptability and specialists playing specific key roles in maintaining community function (Sun et al. 2024b, Zhou et al. 2024). Similarly, abundant and rare taxa from the same community also play important but distinct roles in microbial community function. Abundant taxa usually make up most of the community and contribute most of the nutrient cycling and metabolic activity (Pedrós-Alió 2012), but rare taxa maintain critical functional diversity by performing specific functional roles within the community (Jiao et al. 2017, Xu et al. 2021). In marine sediments, abundant taxa have been shown to be primarily shaped by dispersal limitation and are important for community adaptation and stability in the face of environmental stress (Dash et al. 2024, Sun et al. 2024a), while rare taxa were shaped by deterministic processes likely related to competition within the community (Dash et al. 2024). This type of subcommunity analysis is underexplored in reef environments and marine benthic communities, and a recent review of this topic identifies this as an “urgent need” because it is critical for understanding the distribution patterns of microbial communities (Liu et al. 2019).

To address these unknowns, we performed a field study to identify drivers of biofilm bacterial community assembly and biogeography on the reefs of Roatán, Honduras, with the following objectives: (1) characterize bacterial biofilm communities colonizing two critical benthic reef substrates from eleven reef sites, (2) apply an established statistical framework to quantify the relative roles of stochastic and deterministic processes in shaping bacterial communities in the biofilms, and (3) correlate alpha and beta diversity of bacterial communities with a suite of biotic and abiotic environmental parameters to determine the relative influence of each factor in structuring communities. The latter two objectives were performed for whole communities and within subcommunities of rare and abundant bacterial taxa and of habitat generalists and specialists to look for differential patterns of structure. In this study, we found limited spatial structuring of biofilm communities over 11 km. Instead, the most important factor shaping biofilm bacterial community structure was substrate, as biofilms from different substrates demonstrated distinct diversity, composition, and interactions with environmental conditions. Biofilm bacterial community assembly was dominated by stochastic processes, primarily drift, and the relative influence of each assembly process was similar between substrates. The processes driving assembly of subcommunities were distinct between habitat generalists and specialists and between rare and abundant taxa. These results provide novel insight into the factors shaping biofilm bacterial community assembly and biogeography on two key reef substrates.

## 2. MATERIALS & METHODS

### 2.1. Sample collection

Benthic biofilm communities were sampled from eleven sites on the northwestern coast of Roatán, Honduras between 16 July and 3 August 2023 (Figure 1). Sites were selected to reflect a breadth of environmental conditions and a range of geographic distances in consultation with collaborators from the Roatán Institute for Marine Sciences who are very familiar with the reefs in this area. At each site, biofilm communities were sampled in triplicate at 9 ± 1.5 m depth for two different reef substrates, crustose coralline algae (CCA) and dead coral skeletons, hereby called “calcium carbonate substrate”. Detailed sampling methods can be found in Supplemental Text S1. Briefly, 1.6 cm-diameter cores were removed from the benthos and swabbed using a high-retention microbial swab (Puritan Medical Products, Guilford, ME). Swabs were stored in DNA/RNA Shield (Zymo Research, Irvine, CA) at –20°C prior to processing. Additionally, at each site, a 1-L seawater sample was collected at 9-m depth for nutrient and microbial community analysis. Approximately 40 mL of each water sample was immediately frozen at – 20°C for nutrient analysis. The remaining volume from each water sample was syringe-filtered through a 0.22-µm Sterivex filter (MilliporeSigma, Rockville, MD) and stored in DNA/RNA Shield to capture the bacterioplankton communities.

**Figure 1.**
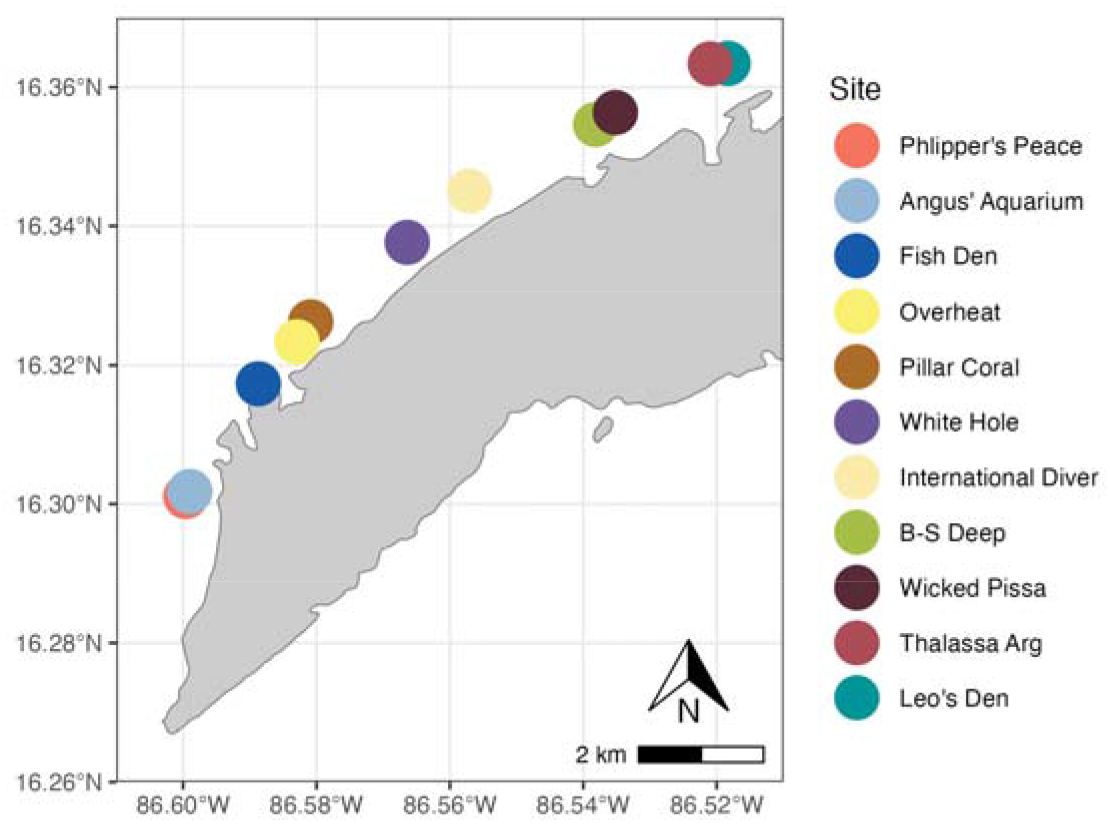
Map of the western point of Roatán, Honduras with points representing the eleven reef sites.

At each site, geographic coordinates and site-level environmental characteristics were recorded during sampling. Temperature was recorded as image metadata by the camera used to photograph each sampling location (Olympus Tough TG-7). Sample-level temperature was calculated by averaging the temperature metadata of approximately five photos taken directly above each sampling location, and the mean and standard deviation of the temperature metadata of all photos from each site (∼30 photos) were used as site-level temperature metrics. An additional 100-mL water sample was collected at 9-m depth for measurement of pH and salinity (Apera Instruments, PH20; Tekcoplus Ltd., SMTK-50). Photographs were taken for three replicate 10-m phototransects at each site to collect site-level coral and algae cover and composition data. Site-level environmental parameters, including coral and macroalgae diversity metrics and concentration of ammonium, combined nitrate and nitrite, and silicate were processed as described in Supplemental Text S2.

### 2.2. DNA extraction, sequencing, and bioinformatics

Genomic DNA was extracted from the swab and filter samples using QIAGEN DNeasy PowerBiofilm Kits (Qiagen, Germantown, MD) following the manufacturer’s protocol with the modifications described in Supplemental Text S3. Genomic DNA from all samples and a DNA extraction kit blank was sent to the Integrated Microbiome Resource (Dalhousie University, Halifax, Nova Scotia, Canada) for amplicon sequencing of the full length 16S small subunit rRNA gene of bacteria and archaea on a PacBio Sequel2 system as described in Comeau (2022) using the forward primer 27F (5’-AGRGTTYGATYMTGGCTCAG-3’) and the reverse primer 1492R (5’-RGYTACCTTGTTACGACTT-3’). Briefly, raw sequences were filtered and trimmed, dereplicated, clustered into operational taxonomic units (OTUs) using a 99% identity cut-off, and chimeric sequences were removed using DADA2 (version 1.26.0, Callahan et al. 2016) and QIIME2 (version 2024.10, Bolyen et al. 2019, QIIME2 Development Team 2023). Taxonomy was assigned using the SILVA 138 reference database and aligned to create a phylogenetic tree (Quast et al. 2013). Detailed bioinformatic methods are provided in Supplemental Text S4.

All sequence data were analyzed in RStudio using the phyloseq (version 1.42.0), tidyverse (version 2.0.0), and vegan (version 2.6.4) packages (McMurdie & Holmes 2013, Wickham et al. 2019, Oksanen et al. 2022). Sequences that were unassigned, identified as mitochondrial or chloroplast DNA, or present in the negative control were removed from the dataset. The bacterial dataset was then split into two parts: biofilm bacterial samples, hereby referred to as “biofilm communities”, and bacterioplankton samples. Sequencing of two carbonate-associated biofilm samples failed, producing <3 reads each after filtering, so these samples were removed from analysis.

### 2.3. Analysis of environmental conditions among sites

Pairs of environmental covariates with correlations with an absolute value >0.7 were considered collinear, and one of the covariates from each pair was removed from analysis. Based on this criterion, the macroalgae Simpson diversity, coral species richness, and coral Shannon diversity parameters were removed from analysis, whereas average sample temperature, average site temperature, depth, salinity, pH, ammonium, combined nitrate and nitrite, silicate, macroalgae cover and Shannon diversity, and coral cover and Simpson diversity were independent and thus retained. Mantel tests with 9,999 permutations were used to assess the correlation between environmental conditions and geographic distance among sites. Prior to the test, the environmental parameters were scaled and centered, and environmental distance was calculated based on Euclidean distances. Coral and macroalgae community distances were calculated using Bray-Curtis distances on Hellinger-transformed community matrices. Geographic distances were calculated from latitude and longitude using Haversine distances.

Bacterioplankton community diversity and composition were characterized for each site. For characterization of alpha diversity, including Shannon-Wiener and Simpson’s diversity indices, bacterioplankton communities were rarefied to the lowest sequence read depth at 7,277 reads per sample. Characterizations of bacterioplankton community composition among sites were performed on unrarefied data by calculating weighted and unweighted UniFrac distances and visualizing with non-metric multidimensional scaling (NMDS) plots (Lozupone & Knight 2005). Finally, to assess correlations between the bacterioplankton community composition and both environmental conditions and geographic distances among sites, mantel tests with 9,999 permutations were used. Environmental and geographic distances were calculated as previously described. Genetic distances among bacterioplankton communities were calculated with weighted and unweighted UniFrac distance metrics based on the Hellinger-transformed OTU table (Legendre & Gallagher 2001).

### 2.4. Analysis of bacterial communities among sample types

All comparisons of alpha diversity among sample types, including bacterioplankton, CCA-associated biofilm, and carbonate-associated biofilm communities, were performed on rarefied datasets. Read depth was determined based on rarefaction curves, and for comparisons of alpha diversity among sample types, the data were rarefied to 2,939 reads per sample. Samples with lower read depth were removed from this analysis. Due to extremely high community diversity, rarefaction curves based on OTU richness did not level off for any sample at any read depth (Supp. Fig. S1a), so OTU richness was not used for any comparisons of alpha diversity. However, both Shannon-Wiener and Simpson’s diversity indices leveled off in all samples (Supp. Fig. S1b-c), so these two metrics were used for comparisons of alpha diversity.

Alpha diversity metrics, including Shannon and Simpson diversity indices, were calculated based on the rarefied communities and compared among sample types and sites using analysis of variance (ANOVA) or Kruskal-Wallis tests, depending on the assumptions of each test. Post hoc testing was conducted for significant results using Tukey’s tests or multiple Mann-Whitney U-tests with Bonferroni correction (Dunn 1961), respectively. To assess differences in bacterial community composition and dispersion among sample types, pairwise distances between samples were calculated using weighted and unweighted UniFrac distances on the unrarefied datasets and compared using permutational multivariate analysis of variance (PERMANOVA) tests with 9,999 permutations and permutation tests for homogeneity in multivariate dispersion (PERMDISP) tests with 9,999 permutations. Post hoc testing was conducted for significant results using the pairwiseAdonis package (version 0.4, Martinez Arbizu 2020).

### 2.5. Analysis of CCA- and carbonate-associated biofilm communities among sites

Due to the significant differences observed in biofilm communities from different substrates, the biofilm sequence dataset was subsetted into separate CCA- and carbonate-associated biofilm datasets for all subsequent analyses. For comparisons of biofilm community alpha diversity among sites, the CCA- and carbonate-associated biofilm datasets were rarefied to 4,684 and 2,939 reads per sample, respectively. All samples with fewer reads than the rarefied depth were removed from alpha diversity analyses. Alpha diversity metrics were calculated and compared among sites using ANOVA or Kruskal-Wallis tests, and differences in community composition and dispersion among sites for CCA- and carbonate-associated biofilm communities were assessed with PERMANOVA and PERMDISP tests, as described previously. To assess correlations between the biofilm community composition and geographic distance among sites, mantel tests with 9,999 permutations were used. Geographic distances were calculated as previously described. Genetic distances among CCA- and carbonate-associated biofilm communities were calculated with weighted and unweighted UniFrac distance metrics based on the Hellinger-transformed OTU tables.

Distance-based redundancy analysis (db-RDA) was used to investigate the influence of individual environmental parameters on biofilm composition. Prior to all db-RDA tests, read counts were transformed using the Hellinger method, and all continuous covariates were centered and scaled. Distances were calculated using weighted and unweighted UniFrac metrics. The best model was chosen using backward selection, where non-significant terms with the lowest explained variation were removed from the full model sequentially until all terms were significant. Of the produced models, the model with the highest adjusted R2 value was selected as the best model. The significance of the best model and each factor in the best model were determined using a permutation test with 9,999 permutations.

### 2.6. Analysis of processes structuring community and sub-community assembly

To investigate the relative influences of selection, dispersal, and drift on the biofilm communities, an analysis was performed with the program Infer Community Assembly Mechanisms by Phylogenetic bin-based null model analysis (“iCAMP analysis”), developed by Ning et al. (2020) based on a previous statistical framework (Stegen et al. 2013), implemented using the iCAMP package (version 1.5.12). Model parameters used are provided in Supplemental Text S5. The relative influence of each ecological process within each bin was then weighted by the relative abundance of the bin to determine the relative influence of each ecological process on the whole biofilm community. Significant differences in the relative influence of each ecological process between substrates were determined by bootstrapping with 1000 repetitions.

Because the processes shaping microbial assembly may vary between whole communities and sub-communities, bins from the iCAMP analysis were then assigned to sub-communities of habitat generalists or specialists and abundant or rare taxa, and the relative influence of each process was then re-calculated at the sub-community-level to compare between the whole community and the sub-community. The EcolUtils package (version 0.1, Salazar 2025) was used to calculate the niche width of each bin with Levins’ index (Levins 1968) and compare the value to a null distribution generated by 9,999 permutations of the OTU table. Bins with significantly high indices were more widely distributed than expected based on their abundance and were classified as habitat generalists, while bins with significantly low indices were narrowly distributed and classified as habitat specialists (Xu et al. 2020, 2022, Isabwe et al. 2022). Bins were also assigned as abundant or rare taxa according to existing relative abundance thresholds, where bins present at <0.001% abundance in the whole community are rare and those present at >0.1% abundance are abundant (Logares et al. 2014, Liu et al. 2015, Mo et al. 2018).

### 2.7. Analysis of environmental factors driving assembly processes in biofilm communities

The role of environmental characteristics, including seawater parameters, bacterioplankton community composition, and nearby coral and algae community composition and diversity, in shaping CCA- and carbonate-associated biofilm composition was investigated on the whole community-level and on phylogenetic bins from the iCAMP analysis that were associated with different assembly processes. To assess whole community patterns, mantel tests with 9,999 permutations were used to correlate biofilm community composition with environmental conditions. Environmental parameters were scaled and centered, and environmental distance was calculated based on Euclidean distances. Genetic distances among CCA- and carbonate-associated biofilm communities were calculated with weighted and unweighted UniFrac distance metrics based on the Hellinger-transformed OTU tables. Mantel tests with 9,999 permutations were used to assess correlations between biofilm community composition and nearby bacterioplankton, coral, and macroalgae community composition.

Bacterioplankton genetic distances were calculated as previously described. Coral and macroalgae community distances were calculated using Bray-Curtis distances on Hellinger-transformed community matrices. Phylogenetic bins were then sorted by their dominant process, defined as the assembly process with the highest relative influence in each bin. Mantel tests with 9,999 permutations were used to assess correlations between communities of selection- and dispersal-dominated bins and nearby bacterioplankton, coral, and macroalgae community composition, as described previously. To determine the phylogenetic bins with the strongest relationship with the measured environmental parameters, generalized linear models with a negative binomial distribution were fit to the abundance of each rarefied taxon with the package MaAsLin2 (version 1.10.0, Mallick et al. 2021). The measured environmental parameters were included as fixed effects.

## 3. RESULTS

### 3.1. Environmental conditions among sites

Environmental conditions were variable among sites, with no clear spatial trends (Supp. Fig. S2). Additionally, differences in environmental conditions, benthic composition, and bacterioplankton community composition were not significantly correlated with geographic distance (Mantel tests, p > 0.209), except for coral community composition, where sites that are closer together have more similar coral communities (Mantel test, r = 0.263, p = 0.021). There was also no significant correlation between bacterioplankton community composition and environmental conditions using weighted (Mantel test, p = 0.468) or unweighted (p = 0.697) UniFrac distances. Detailed descriptions of site-level environmental characteristics are provided in Supplemental Text S6.

### 3.2. Bacterial communities among sample types

There were 2,891,846 raw reads recovered by amplicon sequencing of the 16S rRNA gene of 66 biofilm samples. After quality filtering, 1,647,557 reads remained, corresponding to 162,801 OTUs. OTUs that were identified as mitochondrial (n = 20,003) or chloroplast (n = 12,362) DNA and those found in the negative control (n = 454) were removed. After all filtering steps, 1,136,428 reads corresponding to 129,982 OTUs remained in the final dataset for analysis. The carbonate- and CCA-associated biofilms contained bacterial communities characterized by diverse taxa, including the phyla Bacteroidota, Cyanobacteria, Planctomycetota, Pseudomonadota (both Alpha- and Gammaproteobacteria), and Verrucomicrobiota (Figure 2).

**Figure 2.**
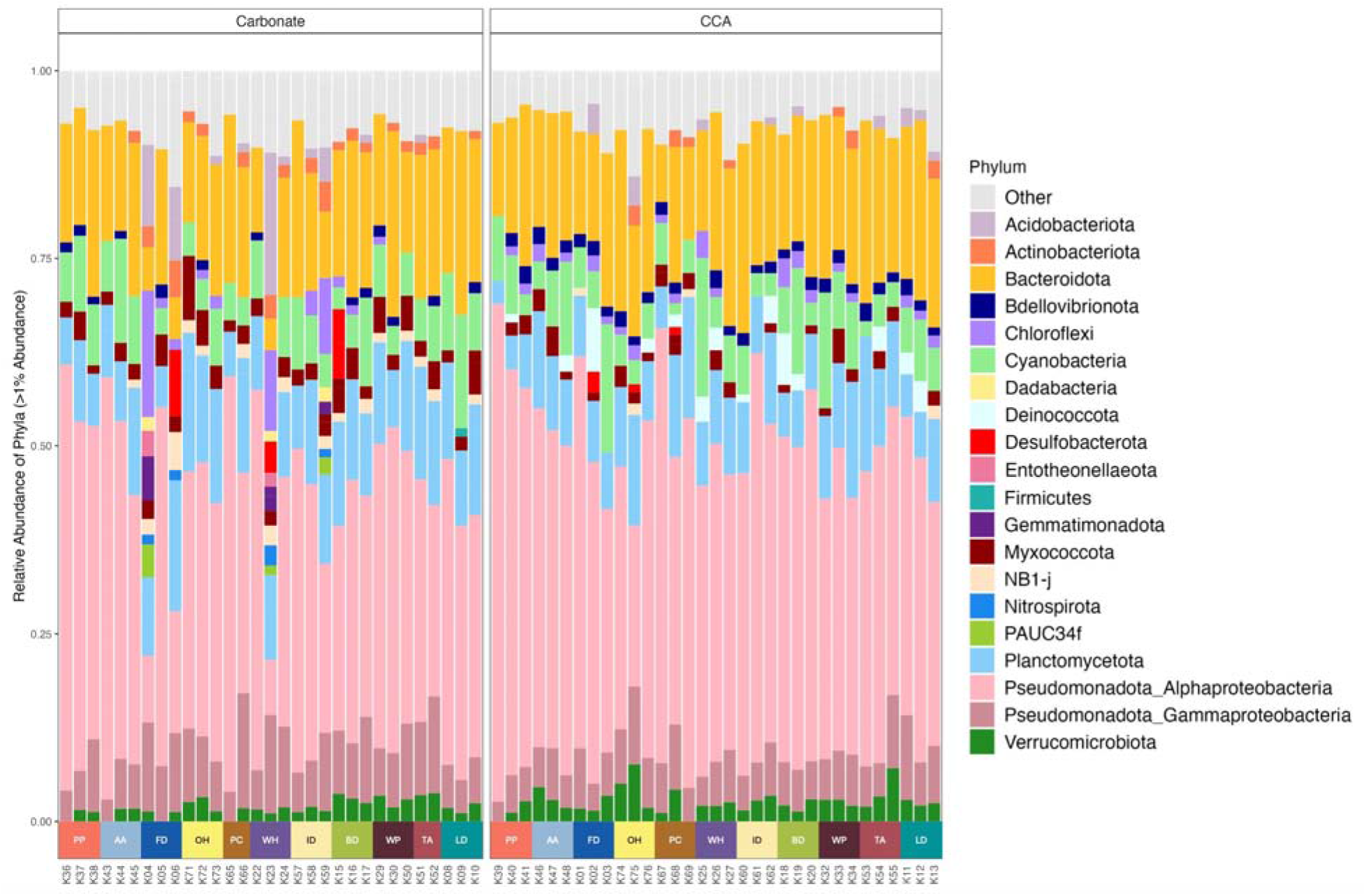
Relative abundance taxa bar plot showing diverse and variable bacterial community composition of carbonate- and crustose coralline algae- (CCA-)associated biofilms. Each bar represents one sample, colored by phylum, and the samples are grouped within each substrate by site. The “Other” group represents all taxa present at <1% abundance in each sample. Site codes: Phlipper’s Peace (PP), Angus’ Aquarium (AA), Fish Den (FD), Overheat (OH), Pillar Coral (PC), White Hole (WH), International Diver (ID), B-S Deep (BD), Wicked Pissa (WP), Thalassa Arg (TA), and Leo’s Den (LD). Sites are ordered by geography from west to east.

Carbonate-associated biofilm communities had significantly higher alpha diversity than CCA-associated biofilm communities (Shannon: t-test, p = 0.001; Simpson: Wilcoxon test, p = 0.002), and biofilm communities on both substrates had significantly higher Shannon and Simpson diversity than bacterioplankton communities (Wilcoxon tests, p < 0.001 for all, Figure 3a). Community dispersion was similar in carbonate- and CCA-associated biofilm communities based on weighted and unweighted UniFrac distances (PERMDISP, weighted: p = 0.16, unweighted: p = 0.256). Dispersion in biofilm communities from both substrates was significantly higher than in bacterioplankton communities according to both metrics (PERMDISP, p < 0.033 for all). Community composition was significantly different among all three sample types according to weighted and unweighted UniFrac distances (PERMANOVA, p = 0.001 for all, Figure 3b). Biofilm community composition was weakly correlated with environmental conditions in CCA-associated communities (Mantel test, unweighted: r = 0.168, p = 0.010, weighted: r = 0.127, p = 0.044) but not in carbonate-associated communities (unweighted: p = 0.715, weighted: p = 0.528).

**Figure 3.**
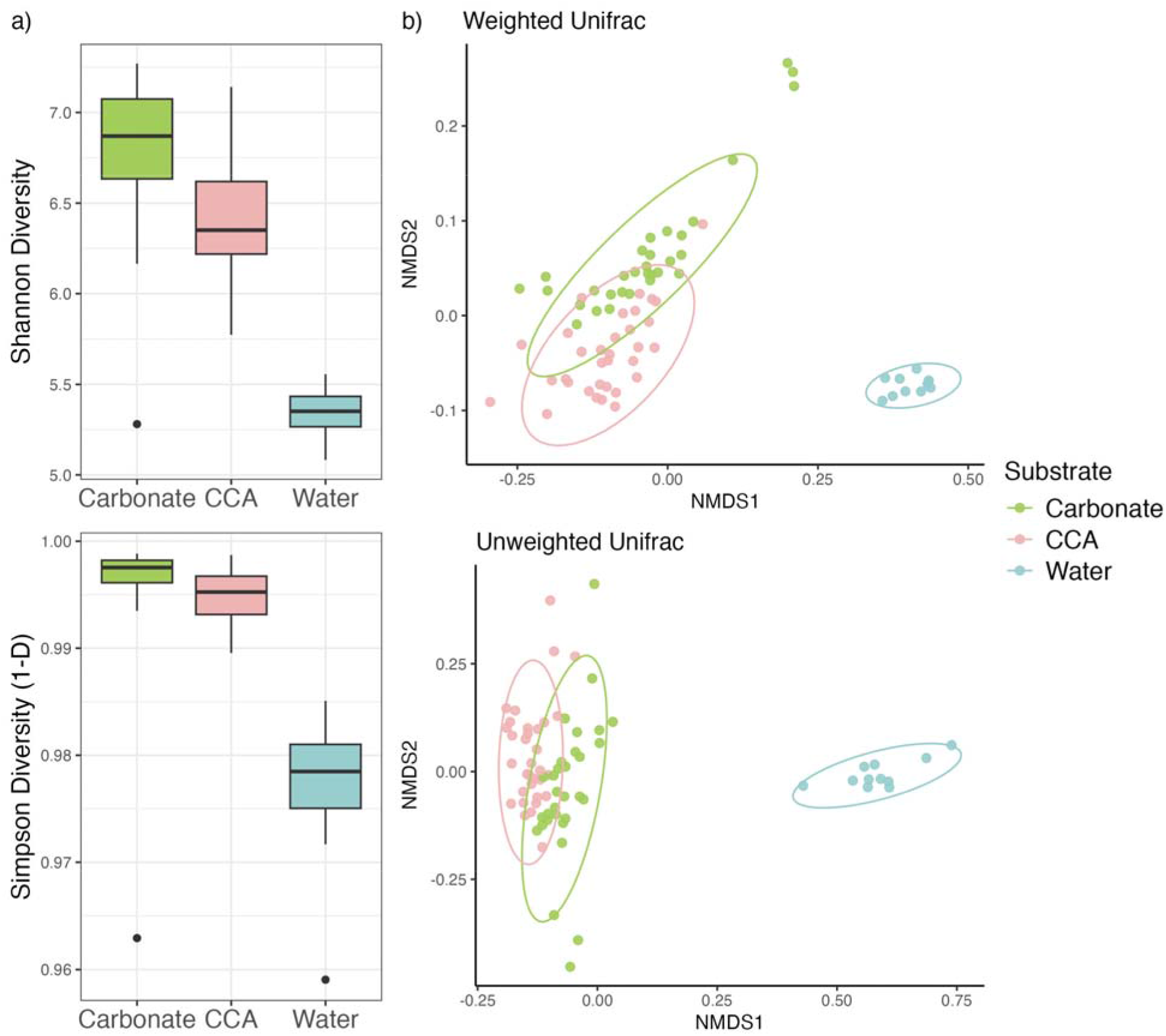
**a)** Alpha diversity and **b)** community composition of carbonate-associated and crustose coralline algae- (CCA-)associated biofilms and bacterioplankton communities. Simpson diversity is presented as 1–D. For NMDS plots, distances were calculated based on weighted and unweighted UniFrac metrics.

Distance-based RDA was used to investigate which environmental variables were significantly associated with changes in biofilm community composition in carbonate- and CCA-associated communities separately. For the CCA-associated communities, the best db-RDA model using weighted UniFrac distances included pH, silicate, and nearby coral cover as covariates, and silicate was weakly correlated but significant (adjusted R^2^ = 0.119, model p < 0.001, p = 0.004 for silicate, permutation test). The best db-RDA model using unweighted UniFrac distances included was the full model, but only the contribution from silicate was significant on its own (adjusted R^2^ = 0.035, model p = 0.034, p = 0.013 for silicate, permutation test). For the carbonate-associated communities, the best db-RDA model based on weighted UniFrac distances included average site temperature, salinity, pH, combined nitrate and nitrite, and silicate as covariates, but only combined nitrate and nitrite concentrations were significantly weakly correlated (adjusted R^2^ = 0.113, model p = 0.028, p = 0.007 for combined nitrate and nitrite, permutation test). The best db-RDA model for carbonate-associated communities based on unweighted UniFrac distances was not significant (p = 0.156, permutation test).

### 3.3. CCA- and carbonate-associated biofilm communities among sites

There were no significant differences in alpha diversity, beta dispersion, or community composition in CCA- or carbonate-associated biofilm communities among sites according to any assessed metric. Additionally, neither CCA- nor carbonate-associated biofilm community composition were significantly correlated with geographic distance between samples based on weighted or unweighted UniFrac distances (Mantel tests, p > 0.223 for all). Details of these results are provided in Supplemental Text S7.

### 3.4. Processes structuring biofilm community and sub-community assembly

Based on the iCAMP analysis, the relative influences of assembly processes contributing to biofilm community composition were similar on carbonate and CCA substrates, despite differences in composition and diversity between the substrates. In biofilms from both substrates, drift (DR) dominated, followed by dispersal limitation (DL), homogeneous (HoS) and heterogeneous (HeS) selection, and finally low levels of homogenizing dispersal (HD, Figure 4). The only significant difference between substrates was the significantly higher influence of homogenizing dispersal in CCA-associated biofilm communities compared to carbonate-associated communities (bootstrapping test, p = 0.012).

**Figure 4.**
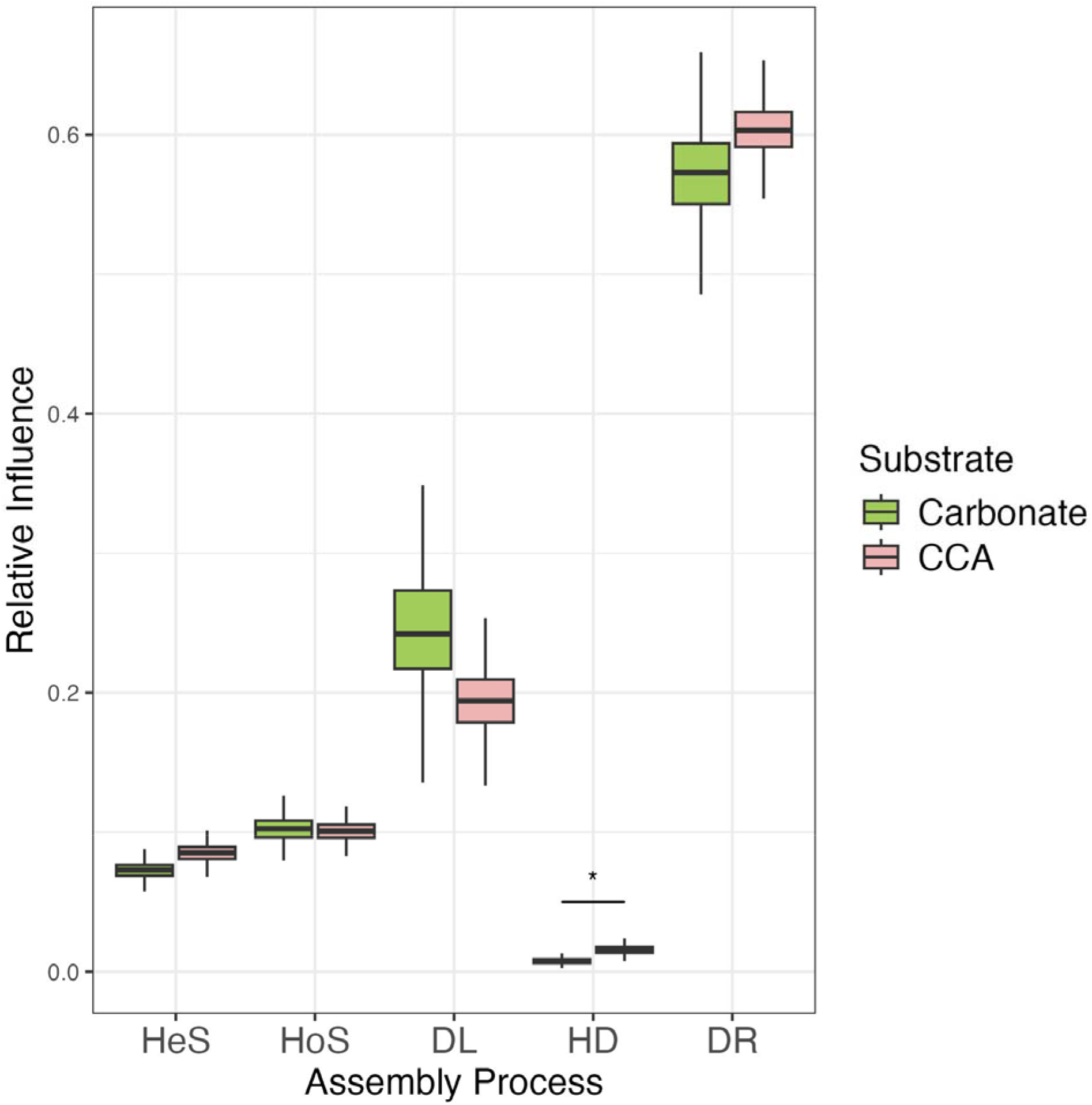
Relative influence of assembly processes in shaping crustose coralline algae- (CCA-) and carbonate-associated biofilm communities. Boxplots represent distributions based on 1000 bootstrapped replicates. HeS = heterogeneous selection, HoS = homogeneous selection, DL = dispersal limitation, HD = homogenizing dispersal, DR = drift. * represents p = 0.01.

To test whether the relative influence of processes structuring sub-communities of habitat generalists and specialists and of rare and abundant taxa match that of the whole communities, the bacterial genera present in the biofilm communities were assigned to sub-communities based on their abundance and distribution. Of the 1,234 genera, 43 were habitat generalists (3.61% abundance), 548 were habitat specialists (68.7% abundance), 155 were abundant (89.1% abundance), and 509 were rare (0.16% abundance, Figure 5a–b). The relative influences of assembly processes differed between sub-communities of habitat generalists and specialists and between abundant and rare taxa in both carbonate- and CCA-associated biofilm communities (Figure 6). Habitat specialists and abundant taxa generally reflected the same influences from assembly processes as the whole communities. Habitat generalists demonstrated higher influence from HeS and HoS and reduced influence of DL compared to specialists, and rare taxa exhibited reduced influence from HeS and HoS and higher influence of DR compared to abundant taxa.

**Figure 5.**
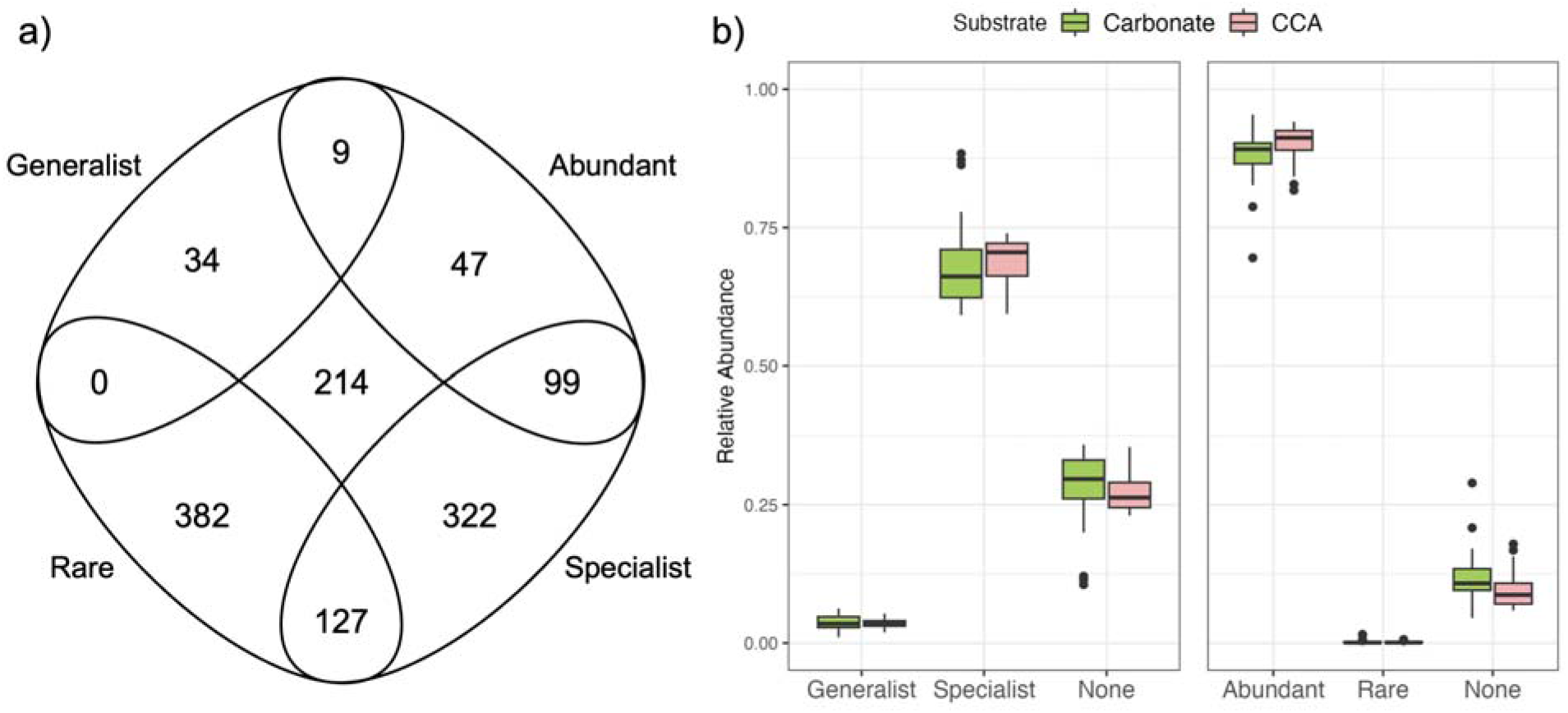
Genus richness and relative abundance of subcommunity members. **a)** Venn diagram showing the number of genera assigned to each sub-community. Two hundred fourteen genera were not assigned to any sub-community. **b)** Boxplots showing the relative abundance of each subcommunity within the whole crustose coralline algae- (CCA-) or carbonate-associated communities by substrate.

**Figure 6.**
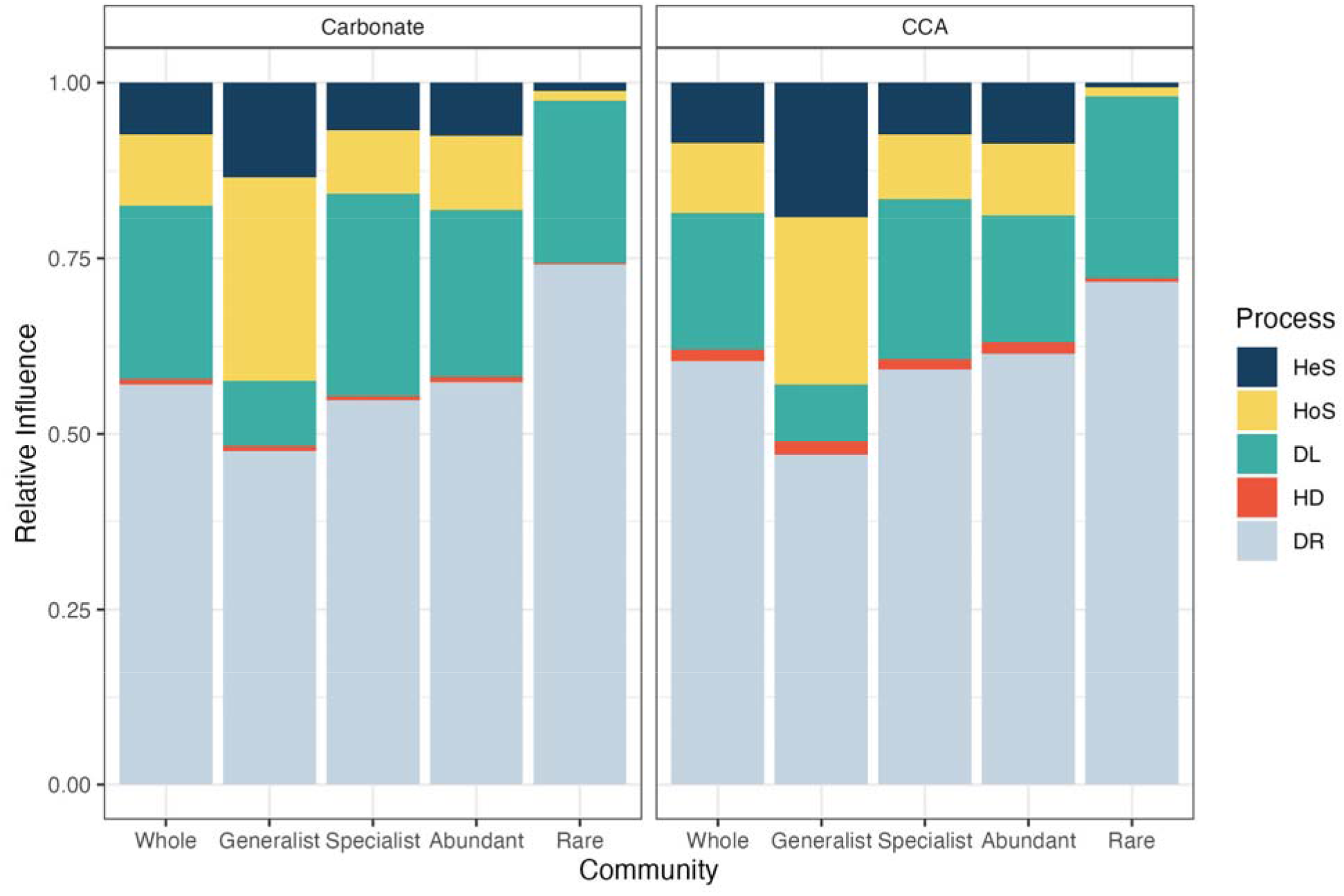
Relative influence of assembly processes in shaping whole biofilm crustose coralline algae- (CCA-) and carbonate-associated communities and sub-communities of habitat generalists, habitat specialists, abundant taxa, and rare taxa. Panels are separated by substrate. HeS = heterogeneous selection, HoS = homogeneous selection, DL = dispersal limitation, HD = homogenizing dispersal, DR = drift.

### 3.5. Environmental factors driving assembly processes in biofilm communities

Mantel tests based on weighted and unweighted UniFrac distances were performed to assess the relationship between nearby communities of bacterioplankton, coral, and macroalgae and both whole biofilm communities and the portions of communities primarily driven by selection and dispersal. The composition of CCA-associated biofilm communities showed no significant correlations with nearby bacterioplankton, coral, or macroalgae community composition, including whole biofilm communities and the community members assembled primarily by selection and dispersal (p > 0.057 for all), except for selection-assembled sub-communities, which were significantly correlated with nearby coral composition based on weighted (r = 0.182, p = 0.034) and unweighted (r = 0.345, p < 0.001) UniFrac distances. In contrast, the composition of whole carbonate-associated biofilm communities was significantly correlated with bacterioplankton community composition based on unweighted UniFrac distances only (r = 0.222, p = 0.023). When the carbonate-associated biofilm community was divided into members primarily assembled by selection and those primarily assembled by dispersal, the selection-assembled sub-community was not significantly correlated with bacterioplankton communities (p = 0.367), but the dispersal-assembled sub-community was (weighted: r = 0.200, p = 0.024). The only other significant comparison was the correlation between nearby coral community composition and carbonate-associated biofilm community members primarily assembled by dispersal (weighted UniFrac, r = 0.182, p = 0.052). All p- values associated with these Mantel tests are reported in Table 1.

**Table 1.** P-values associated with Mantel tests of whole biofilm communities and dispersal- or selection-assembled members of crustose coralline algae- (CCA-) and carbonate-associated biofilm communities with nearby bacterioplankton, coral, and macroalgae communities. “Water” indicates p-values associated with bacterioplankton communities. Significant p-values are marked with *.

|  |  | CCA |  |  | Carbonate |  |  |
| --- | --- | --- | --- | --- | --- | --- | --- |
|  |  | Whole | Dispersal | Selection | Whole | Dispersal | Selection |
| Weighted UniFrac | Water | 0.589 | 0.708 | 0.077 | 0.494 | 0.314 | 0.672 |
|  | Coral | 0.139 | 0.648 | <0.001* | 0.687 | 0.052* | 0.064 |
|  | Macroalgae | 0.389 | 0.165 | 0.783 | 0.793 | 0.840 | 0.999 |
| Unweighted UniFrac | Water | 0.275 | 0.616 | 0.073 | 0.023* | 0.024* | 0.367 |
|  | Coral | 0.469 | 0.764 | 0.034* | 0.342 | 0.201 | 0.124 |
|  | Macroalgae | 0.221 | 0.057 | 0.549 | 0.986 | 0.928 | 0.998 |

Finally, to assess the role of environmental conditions in shaping individual assembly processes, the MaAslin2 analysis was used to correlate the suite of environmental parameters with the abundance of each phylogenetic bin produced during the iCAMP analysis, which were then grouped by dominant assembly process. All measured environmental factors were correlated with at least one phylogenetic bin, and the bins with significant correlations with environmental conditions were distinct between substrates (Figure 7). The strongest correlations with environmental conditions were found in bins dominated by DL in communities from both substrates and in bins dominated by HoS from CCA-associated biofilm communities. There were few significant relationships with environmental conditions in bins dominated by selection (i.e., HeS and HoS) from carbonate-associated biofilm communities.

**Figure 7.**
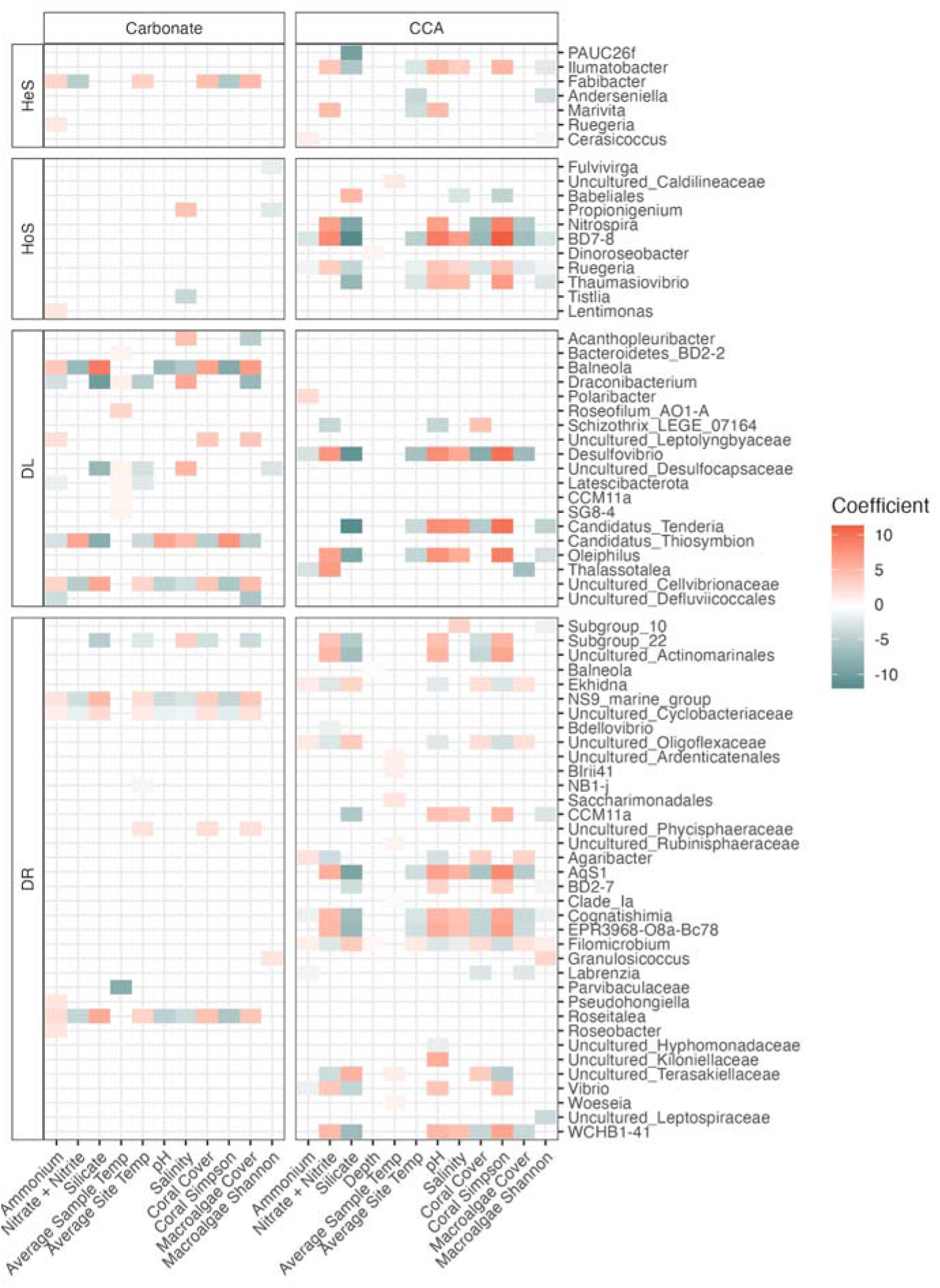
Phylogenetic bins and environmental factors with significant correlations. Each row represents a phylogenetic bin labeled by its dominant genus, which are grouped by the dominant process shaping each bin and the substrate (i.e., carbonate or crustose coralline algae [CCA]). HeS = heterogeneous selection, HoS = homogeneous selection, DL = dispersal limitation, DR = drift. No bins were dominated by homogenizing dispersal. Missing tiles represent non-significant correlations, and bins not represented had no significant correlations with environmental variables.

## 4. DISCUSSION

Benthic biofilm bacteria in reef ecosystems represent an understudied habitat of microbial diversity, characterized by extremely high taxonomic diversity and functional potential distinct from that of the relatively well-described water column (Zhang et al. 2019). Because these biofilms are critical contributors to broader reef health and functioning, understanding the relative influence of ecological and evolutionary assembly processes in these communities and their interactions with environmental conditions is necessary for understanding reef resilience in the face of environmental change. To address these unknowns, we characterized benthic biofilm bacterial communities on two important reef substrates, CCA and carbonate, and quantified the relative contributions of selection, dispersal, and drift in the assembly of those communities.

Despite clear differences in diversity and composition, the biofilms associated with the two substrates showed the same relative contributions from assembly processes, suggesting that microbial community assembly follows consistent patterns independent of microbial taxonomy. Further, we investigated whether the contributions of processes shaping whole communities matched those of sub-communities of 1) habitat generalists and specialists and 2) abundant and rare taxa and found that assembly processes were distinct between sub-communities. These results provide new insight into the relative roles of deterministic and stochastic processes in shaping benthic reef biofilm assembly.

### 4.1. Substrate type, and not site, influences biofilm community structure

Biofilms on neither substrate showed significant differences in bacterial community diversity, dispersion, or composition among the sites, which were separated by a maximum of 11.1 km, and bacterial community composition was not correlated with geographic distance in biofilms associated with either substrate. This indicates that mature biofilms on these two important reef substrates are relatively consistent across the reef system, even in the face of natural environmental heterogeneity (Supp. Fig. S2). While this has been previously observed in biofilms developing in marine environments over small spatial scales (<8 km, Sweet et al. 2011, Tan et al. 2015, Stenger et al. 2024), most studies of reef biofilm biogeography found differences in community composition among sites (5–30 km, Kriwy & Uthicke 2011, Witt et al. 2011, Kegler et al. 2017, Yanovski et al. 2024). However, all of these studies were specifically testing for environmental influences on biofilm composition along directional, previously described water quality gradients stemming from regions of high anthropogenic impacts, so the observed differences were likely driven by these persistent environmental influences.

Together, our results suggest that at this scale, within a single reef system, microbial dispersal is high enough to overcome spatial structuring from dispersal limitation, with differences in biofilm community structure primarily shaped by local factors, in line with the Baas Becking hypothesis (Hanson et al. 2012, Ragon et al. 2012). This aligns with the general consensus that the relative influence of assembly processes varies depending on the scale of analysis, with local scales (<10 km) primarily driven by environmental filtering and the impact of dispersal limitation increasing with geographic scale (Martiny et al. 2006, Liu et al. 2023a, Xu et al. 2024, Soininen & Graco Roza 2024). This is the first time this pattern has been characterized in climax biofilm communities from natural reef substrates, but future studies should expand on this work to elucidate patterns of assembly in natural climax communities across spatial scales to confirm the generalizability of this finding. If the impacts of microbial dispersal limitation are negligible and local factors dominate microbial assembly over small spatial scales, this supports the use of microbial bioindicators in reef monitoring schemes, has implications for site selection during biofilm-based coral restoration methods, and suggests that reef biofilms may be sensitive to future environmental shifts that could cause cascading impacts on ecosystem health and function.

In the absence of strong spatial drivers, the dominant local factor shaping these communities was substrate type, evidenced by significant differences in composition and diversity between carbonate- and CCA-associated biofilm communities. This suggests that substrate heterogeneity on reefs is an important driver of ecosystem-level microbial diversity. As reef biofilms represent a potentially enormous source of unexplored taxonomic diversity on our planet (Sneed et al. 2015, Galand et al. 2023, Hochart et al. 2024), more work needs to be done to understand the regulation and assembly of biofilm communities across diverse substrates to better elucidate the diversity and function of reef biofilms (Dang & Lovell 2015). Biofilms have commonly been studied on artificial substrates and in early successional stages, but studies of climax communities on natural reef substrates, such as this one, are lacking. Despite this, the results of this study align with the previous literature, which found significant differences in bacterial community composition across a variety of natural and artificial substrates (Sweet et al. 2011, Witt et al. 2011, Sajid et al. 2024). This finding is typically attributed to differences in surface properties of the substrate, such as texture and shape (Sweet et al. 2011, Dang & Lovell 2015). Interestingly, despite significant differences in community composition between CCA- and carbonate-associated communities, the dominant phyla in biofilms from both substrates were similar and matched those described in the literature for diverse reef substrates, including Alphaproteobacteria, Bacteroidota, Planctomycetota, Cyanobacteria, and Verrucomicrobiota (Kriwy & Uthicke 2011, Kegler et al. 2017, Yang et al. 2021, Yanovski et al. 2024, Sajid et al. 2024, Hochart et al. 2024).

CCA- and carbonate-associated biofilm communities additionally differed in their relationship to environmental conditions. CCA-associated biofilm community composition was significantly correlated with environmental conditions, particularly silicate concentrations, while carbonate-associated community composition was not. Surface biofilms of CCA have been studied extensively due to their connection with coral settlement, and bacterial composition of CCA-associated biofilms is often related to the CCA host species (Sneed et al. 2015, Siboni et al. 2020, Jorissen et al. 2021). This indicates that the algae may be regulating the biofilm present on its surface, as microbial attachment to biotic surfaces is known to be mediated by specific cellular processes from the host (Rendueles & Ghigo 2012, Dang & Lovell 2015). This was observed by Hochart et al. (2024), who demonstrated phylosymbiosis between CCA chlorotype and surface biofilm community composition, suggesting that CCA individuals are selective of their surface microbiomes. This is supported by our finding that whole CCA microbiomes were not influenced by nearby bacterioplankton, coral, or macroalgae communities. In fact, only the selection-assembled portion of CCA-associated bacterial communities was significantly correlated with nearby coral community composition. This could be explained by host regulation of the biofilm composition through the provisioning of essential resources, as this relationship was not observed in carbonate-associated biofilms, and may be the result of tight spatial relationships between the algae, its microbiome, and coral settlers.

The regulation of surface biofilm communities by the host would also explain the lower diversity of CCA-associated biofilms observed compared to carbonate-associated biofilms, though this could also be a product of regular cell sloughing by the algae, which is thought to limit the diversity of surface biofilm communities (Sneed et al. 2015). Additionally, bacterial composition of CCA-associated biofilms has previously been linked with environmental conditions (Webster et al. 2011, 2013, 2016, Jorissen et al. 2021), as seen in this study. This suggests a balance between host regulation and environmental drivers in the establishment and maintenance of CCA-associated biofilms (Hester et al. 2016).

Contrary to the CCA-associated biofilms, there are few direct comparisons in the literature to the carbonate-associated biofilms in this study, which are climax communities of surface biofilms from standing dead coral skeletons sampled directly from the reef framework. However, biofilms developed on autoclaved coral skeletons and sterile calcium carbonate for 2– 7 weeks resembled those in this study with high diversity and stability across samples (Witt et al. 2011, Sajid et al. 2024). Kegler et al. (2017) sampled climax biofilm communities from small rocks with some CCA found on the reef and found the communities to be significantly correlated with combined nitrate and nitrite, like the carbonate-associated biofilms in this study, and silicate, like the CCA-associated biofilms in this study, as well as phosphate, chlorophyll *a*, and suspended particulate matter, which were not assessed in this study. This suggests that carbonate-associated biofilms may have weaker associations with environmental conditions than CCA-associated biofilms or are correlated with parameters not measured in this study.

Because carbonate is an abiotic surface, carbonate-associated biofilms are not limited by host regulation and are more likely to be related to surface characteristics such as surface charge and hydrophobicity (Rendueles & Ghigo 2012). However, results from this study also showed that carbonate-associated biofilms, and especially the dispersal-assembled members of those biofilms, have significant correlations with nearby bacterioplankton and benthic community composition, indicating that interactions with nearby organisms play some role in the movement of bacterial taxa across the reef landscape and in structuring carbonate-associated biofilm communities that was not observed for CCA-associated communities. It is possible that carbonate-associated biofilms are regulated by the turf algae that also colonize the carbonate surface. Communities of turf algae thrive in nutrient-rich environments and are known to promote the proliferation of copiotrophic and pathogenic bacteria through the release of nutrient-dense exudates (Vermeij et al. 2009, Nelson et al. 2013, Sweet et al. 2013). Regulation of carbonate-associated biofilm composition by nearby turf algae could explain why only db-RDA models based on weighted UniFrac distances were significant for carbonate-associated biofilm communities, as the turf algae moderate the abundance but not presence of certain bacterial taxa. Taken together, these findings highlight key differences in biofilm community structure between two critical reef substrates and emphasize the need for further investigation of microbial assembly processes across diverse reef substrates.

### 4.2. Stochastic processes dominate biofilm assembly across reef substrates

Despite the significant differences in biofilm diversity, composition, and relationship to environmental conditions between substrates, the relative contribution of assembly processes to biofilm structure was very similar on both substrates. Drift had the largest relative contribution to biofilm assembly on both substrates (60.4% on CCA, 57.1% on carbonate), followed by dispersal limitation, homogeneous selection, and heterogeneous selection, and the relative contribution of homogenizing dispersal to microbial assembly on both surfaces was extremely low (Figure 4). These results highlight the importance of stochastic processes (i.e., drift and dispersal) to microbial assembly on reefs across key substrates, contradicting the common assumption that large bacterial population sizes and high dispersal rates render drift negligible (Zhou & Ning 2017). The dominance of drift in structuring these communities aligns with the understanding that drift becomes relatively more important in highly productive communities and those drawing from large regional species pools (Chase & Myers 2011).

Given the relatively large influence of dispersal limitation in these communities, it was surprising that no patterns of spatial structure were observed in biofilms associated with either substrate, as dispersal limitation in isolation is known to drive distance-decay relationships in microbial communities (Hanson et al. 2012). This, in combination with the very low relative influence of homogenizing dispersal, suggests that homogeneous selection played a key role in reducing community divergence among sites in this study. Because environmental conditions were also not correlated with geographic distance in this study, the differentiating role of dispersal limitation may have been further balanced by the influence of heterogeneous selection. The results of this study contribute to the understanding that interactions between processes like dispersal and selection may lead to complex or unexpected patterns of assembly (Stegen et al. 2013, Evans et al. 2017, Liu et al. 2019). For example, dispersal limitation relies on drift and diversification to drive community divergence (Nemergut et al. 2013), both of which are included in the “Drift” category as they are indistinguishable by the iCAMP analysis framework (Ning et al. 2020). Future studies should work to incorporate evolutionary models into this type of analysis framework so that diversification can be quantified and the interactive relationships between drift, dispersal, and diversification can be examined, and future experimental studies should be performed to investigate the impacts of assembly processes on communities in isolation and in combination to better understand these interactions.

Bacterial communities in marine environments are often, but not always, shown to be primarily driven by stochastic processes, including in coral microbiomes (Zhang et al. 2021, Zhao et al. 2024), bacterioplankton (Zhang et al. 2021, Ma et al. 2022, Liu et al. 2022, Liang et al. 2024), subtidal and intertidal sediments (Ma et al. 2022, Gong et al. 2022, Li et al. 2023, Cheng et al. 2023, Wang et al. 2023, Xu et al. 2024, Liang et al. 2024), and macroalgae microbiomes (Pearman et al. 2024). However, the magnitude of the influence of stochastic processes observed in this study (81.4% on CCA, 82.5% on carbonate) was only seen in one study of sediments collected from a mangrove forest (Thomson et al. 2022). Bacterial communities in marine systems are known to have relatively high influence from stochastic processes because water currents homogenize environmental conditions and promote long-distance passive dispersal and because marine bacteria have generally evolved to tolerate significant environmental variability (Liu et al. 2019).

There are several possible explanations for the high relative influence of stochastic processes (i.e., drift and dispersal) observed in this study. The extremely high phylogenetic diversity found in reef biofilms, as demonstrated in this study, likely indicates extensive functional redundancy and niche overlap within these communities, which is known to promote drift (Leibold & McPeek 2006, Zhou & Ning 2017). High phylogenetic diversity has also been shown to drive stronger dispersal limitation (Li et al. 2023). Alternatively, deterministic processes (i.e., selection) increase in the face of chronic environmental stress (Chase 2007, Dai et al. 2017, Liu et al. 2023a, Cheng et al. 2023), so the high stochasticity seen in these communities may indicate that environmental stress and anthropogenic impacts are relatively low in the examined region at the time of sample collection (Dini-Andreote et al. 2015, Bahram et al. 2016, Liu et al. 2022). This relationship should be further characterized, as it could serve as a bioindicator for reef managers to track temporal trends in ecosystem-level environmental stress or quantitatively monitor outcomes of implemented water quality management strategies.

Despite major differences in the biofilms from different substrates, the relative contributions of assembly processes were consistent between CCA- and carbonate-associated biofilms for whole communities and both examined classifications of sub-communities, indicating that the underlying factors shaping biofilm community assembly are not substrate-specific but rather characteristics of the environment or the bacteria themselves. It is possible that biofilms on many reef substrates are assembled with similar influence from each process because they experience similar suites of environmental conditions and draw from the same regional species pool, and differences in the biofilm climax community composition are driven by priority effects early in the successional process (Gleason 1927, Nemergut et al. 2013, Debray et al. 2022). Alternatively, dominant assembly processes may be related to specific bacterial traits, such as niche breadth, cell size, biotic interactions, population size, and dormancy (Wu et al. 2018, Aslani et al. 2022), which is supported by the finding that bacteria from subcommunities representing different traits were assembled according to different processes but still assembled similarly between substrates. If this is the case, then the results of this study suggest that CCA and carbonate represent similar niche spaces which are colonized by bacteria with similar traits, even if their composition is distinct. Future studies should continue to investigate assembly processes of biofilms on different substrates to better understand the factors that shape microbial diversity and extend this work to examine functional profiles of bacterial communities from different reef substrates.

### 4.3. Bacterial sub-communities demonstrate distinct assembly patterns

The relative influence of processes contributing to the assembly of some sub-communities was different from those of the whole community on both substrates. In particular, while habitat specialists were structured similarly to the whole communities, habitat generalists were characterized by increased influence of both heterogeneous and homogeneous selection and reduced influence of dispersal limitation. Sub-communities of abundant taxa were structured similarly to the whole communities, but rare taxa experienced less influence from homogeneous and heterogeneous selection and more influence from drift. It is possible that specialists and abundant taxa reflect the same patterns as the whole community because those sub-communities dominate the whole community (Figure 5b), as seen previously (Liao et al. 2016, Li et al. 2023, Nethmini et al. 2025).

The patterns of dispersal limitation and drift observed within the analyzed sub-communities are supported by previous work on marine bacterial assembly. Habitat generalists are characterized by wide habitat tolerance and the ability to exploit diverse resources (Liao et al. 2016) and are regularly seen to have reduced influence of dispersal limitation (Yan et al. 2022, Sun et al. 2024b, Zhou et al. 2024). The results of this study suggest that bacterial generalists in reef biofilms have increased ability to move among patches of suitable habitat in a landscape of highly heterogeneous substrates, relative to specialist taxa. Reduced drift in generalists and increased drift in rare taxa, compared to specialists and abundant taxa, respectively, also align with findings from previous studies (Aslani et al. 2022, Nethmini et al. 2025). The relative influence of drift has been negatively linked with both niche breadth (Hanson et al. 2012) and taxa abundance (Zhou & Ning 2017, Jia et al. 2018, Fodelianakis et al. 2021, Liu et al. 2023b), where narrower niches and lower abundance make taxa more susceptible to the effects of drift. Between these two processes, sub-communities of rare taxa were almost entirely assembled by stochastic processes (98.0% on CCA, 97.5% on carbonate). This pattern suggests that the rare taxa in this study were primarily transient members of the biofilm communities, as strong dispersal limitation means taxa are only occasionally re-introduced and strong drift causes regular local extinctions (Jia et al. 2018).

The impacts of selection on sub-communities of habitat generalists and rare taxa are not so straightforward. The literature generally supports our findings that generalists are structured with higher influence of selection than specialists (Aslani et al. 2022, Yan et al. 2022, Sun et al. 2024b, Nethmini et al. 2025) and rare taxa are structured with lower influence of selection than abundant taxa (Mo et al. 2018, Ji et al. 2020, Liu et al. 2023b, Li et al. 2023, Pearman et al. 2024, Zhang et al. 2025). The strong influence of homogeneous selection in sub-communities of generalists in this study indicates that environmental stability is an important factor in the assembly of generalists, which aligns with previous work that showed that large niche breadth was linked with strong adaptability (Sun et al. 2024a, Zhao et al. 2024). Low homogeneous selection in sub-communities of rare taxa have been shown to encourage biogeographic structuring of rare taxa through significant interactions with environmental parameters and dispersal limitation, while sub-communities of abundant taxa, which had strong influences from selection, demonstrated weak associations with geography (Liu et al. 2023b). This pattern could explain the lack of spatial structuring observed in this study, as abundant taxa dominated the communities and may have swamped any biogeographic signal in the rare taxa. However, several studies of marine bacterial communities have found contrasting results, with higher relative influence of selection in sub-communities of rare taxa (Liu et al. 2015, Dash et al. 2024, Sun et al. 2024a), and a similar lack of consensus has also been noted for non-marine communities (Pearman et al. 2024, Sun et al. 2024a). The discrepancy in the literature suggests that the influence of selection on bacterial sub-communities may be dependent on sample type, environment, or methodology, so future work should consider these factors when designing experiments and developing generalizations in order to accurately assess the disparate patterns of assembly among bacterial sub-communities.

### 4.4. Environmental influence on biofilm assembly is taxon-specific

The impact of environmental conditions on marine biofilm composition has been studied for decades, including the significant differences caused by environmental factors examined in this study, such as inorganic nutrient concentrations (Meyer-Reil & Köster 2000, Chiu et al. 2008, Rao 2010, Sawall et al. 2012, Kegler et al. 2017, Chen et al. 2017, Remple et al. 2021, Cheng et al. 2023), pH (Cheng et al. 2023), salinity (Lau et al. 2005, Chiu et al. 2005, Chen et al. 2017), temperature (White et al. 1991, Lau et al. 2005, Chiu et al. 2005, Rao 2010), depth (Webster et al. 2004), and nearby benthic composition (Sawall et al. 2012, Kegler et al. 2017, Remple et al. 2021), and those not covered in this study, such as tides (Dobretsov & Qian 2006) and heavy metal concentrations (Cheng et al. 2023). Because of the well-established relationship between biofilm composition and many environmental conditions, it was surprising that the compositions of climax communities in this study were correlated with so few environmental parameters. These results demonstrate that patterns of environmental influence on biofilm composition can be inconsistent on the whole community level and are likely context dependent. By splitting the whole communities into phylogenetic bins and grouping them by dominant assembly process, we were able to tease out patterns of environmental influence at the sub- community scale that may explain this apparent inconsistency. This analysis immediately revealed that many more environmental conditions were contributing to community assembly than were discernible at the whole community-scale, as every environmental parameter measured was significantly correlated with at least one phylogenetic bin, except for depth in the carbonate-associated biofilms (Figure 7).

Several emergent patterns contributed to the complexity of these taxa-environment interactions. First, the strength of environmental correlations varied depending on which process dominated the assembly of each bin, with dispersal- and selection-assembled taxa having stronger correlations with environmental conditions than drift-assembled taxa. This indicates that diverse environmental characteristics influence the movement, establishment, and persistence of bacterial taxa in reef biofilms and supports the understanding that dispersal is not a purely stochastic process (Hanson et al. 2012, Nemergut et al. 2013). Second, the patterns of environmental correlations with biofilm taxa differ by substrate. For example, coral Simpson diversity, combined nitrate and nitrite concentrations, and pH had strong positive relationships with taxa in CCA-associated biofilms but primarily neutral or negative relationships with carbonate-associated taxa. Finally, these patterns may be interactive, with environmental conditions structuring the taxa dominated by each assembly process differently in biofilms from different substrates. For example, average sample temperature is most important for dispersal-assembled taxa in carbonate-associated biofilms, but in CCA-assembled biofilms, average sample temperature is mostly related to drift-assembled taxa. Similarly, there are relatively few significant relationships between environmental factors and selection-assembled taxa in carbonate-associated biofilms compared to other substrate-process pairs. Together, these examples highlight that environmental conditions impact bacterial taxa in distinct ways, likely influenced by the traits and ecology of each taxon, and assessing these relationships on a whole community-level may obscure complex patterns, fuel disagreement between studies, and limit understanding of which environmental parameters play significant roles in microbial assembly.

## 5. CONCLUSIONS

Biodiversity is critical to ecosystem functioning, but our understanding of how biodiversity is generated and maintained is lacking, especially in exceptionally diverse reef ecosystems (Zhou & Ning 2017, Liu et al. 2019). Because of this, a central objective in the field of microbial ecology is to characterize the processes driving community assembly and the impacts of spatial and environmental influences on those processes (Martiny et al. 2006, Hanson et al. 2012, Nemergut et al. 2013). To do this, we examined biofilm bacteria colonizing two important reef substrates from sites spanning ∼11 km and found that, on this scale, spatial factors had little influence on community composition or structure. The influences of environmental conditions and substrate type on the biofilm communities, however, suggested that assembly processes were tied to traits of individual bacterial taxa. Whole community analyses are useful for understanding community trends and impacts on greater ecosystem function, but future studies of microbial assembly should prioritize taxon-level analyses, such as the iCAMP model used in this study (Ning et al. 2020), to produce better descriptions of the complex and interactive relationships between bacterial taxa and assembly processes needed to further microbial ecology theory. This takeaway is further supported by the finding that sub-communities of bacterial taxa, including habitat generalists and specialists and rare and abundant taxa, are assembled by distinct patterns of processes that are consistent between substrates.

Overall, the observed dominance of stochastic processes in shaping bacterial assembly in reef biofilms means that predicting shifts in microbial community composition and function in the face of ongoing environmental change will be even more difficult than previously described (Zhou & Ning 2017), so continuing to study the relationships between environmental conditions, individual biofilm-forming taxa, including bacteria and microeukaryotes, and both stochastic and deterministic assembly processes, as well as overall biofilm sensitivity to stochastic assembly, will be critical to characterizing future reef resilience.

## Acknowledgements

We thank Jennifer Keck, Tom Wood, the Roatán Institute for Marine Science’s 2023 Coral Reef Research Interns, and the Anthony’s Key dive masters and boat captains for providing field support. This work was supported by grants from the National Science Foundation Graduate Research Fellowship Program and the American Philosophical Society to JAS. The present work was part of JAS’ PhD dissertation.

## Data availability

The raw 16S rRNA gene amplicon sequencing data generated during this study have been deposited in the NCBI Sequence Read Archive (SRA) under BioProject accession PRJNA1493256. All sample metadata and code required to reproduce the analyses and figures presented in this study are publicly available in the GitHub repository at https://github.com/simsjordan/KARMA.

## Author contributions

JAS designed the experiment, performed field and laboratory work, analyzed the data, and wrote the original manuscript. All authors reviewed manuscript drafts.

## Notes

https://github.com/simsjordan/KARMA

### Competing Interest Statement

The authors have declared no competing interest.

