## Supplemental Figures for "Convergent stochastic assembly governs reef biofilm microbiomes across ecologically distinct benthic substrates"

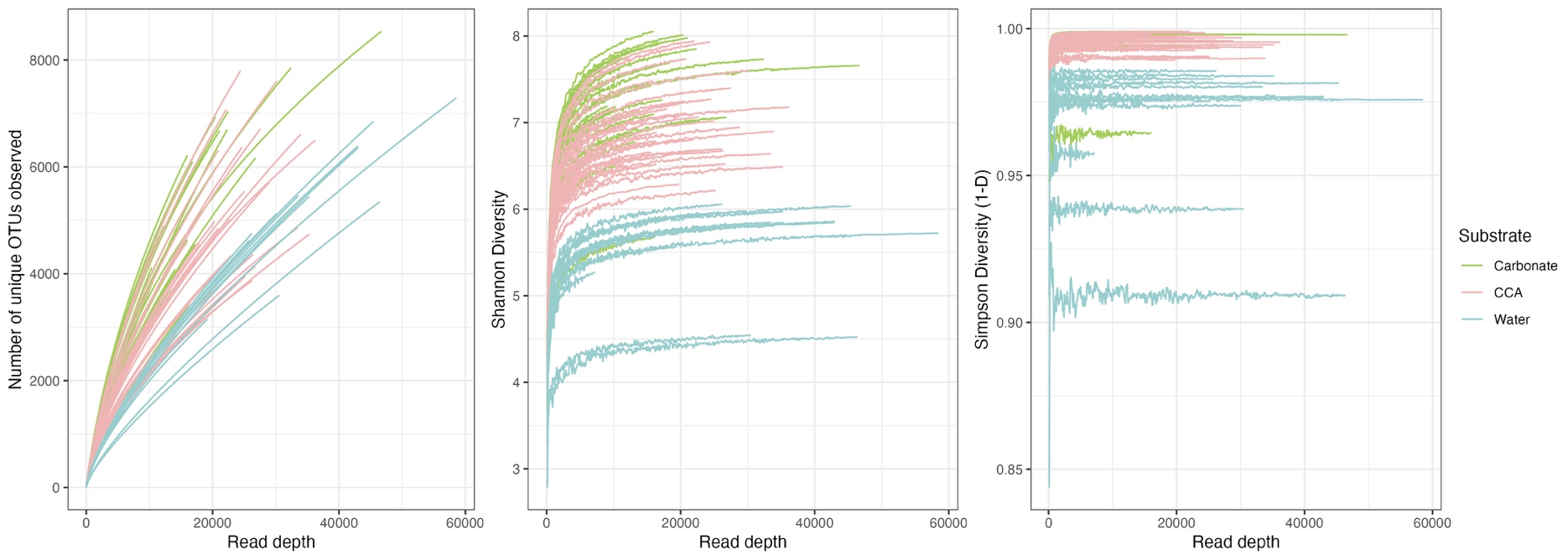
Supplemental Figure S1. Rarefaction curves based on a) OTU richness and b) Shannon and c) Simpson diversity indices. Simpson diversity is presented as 1–D.

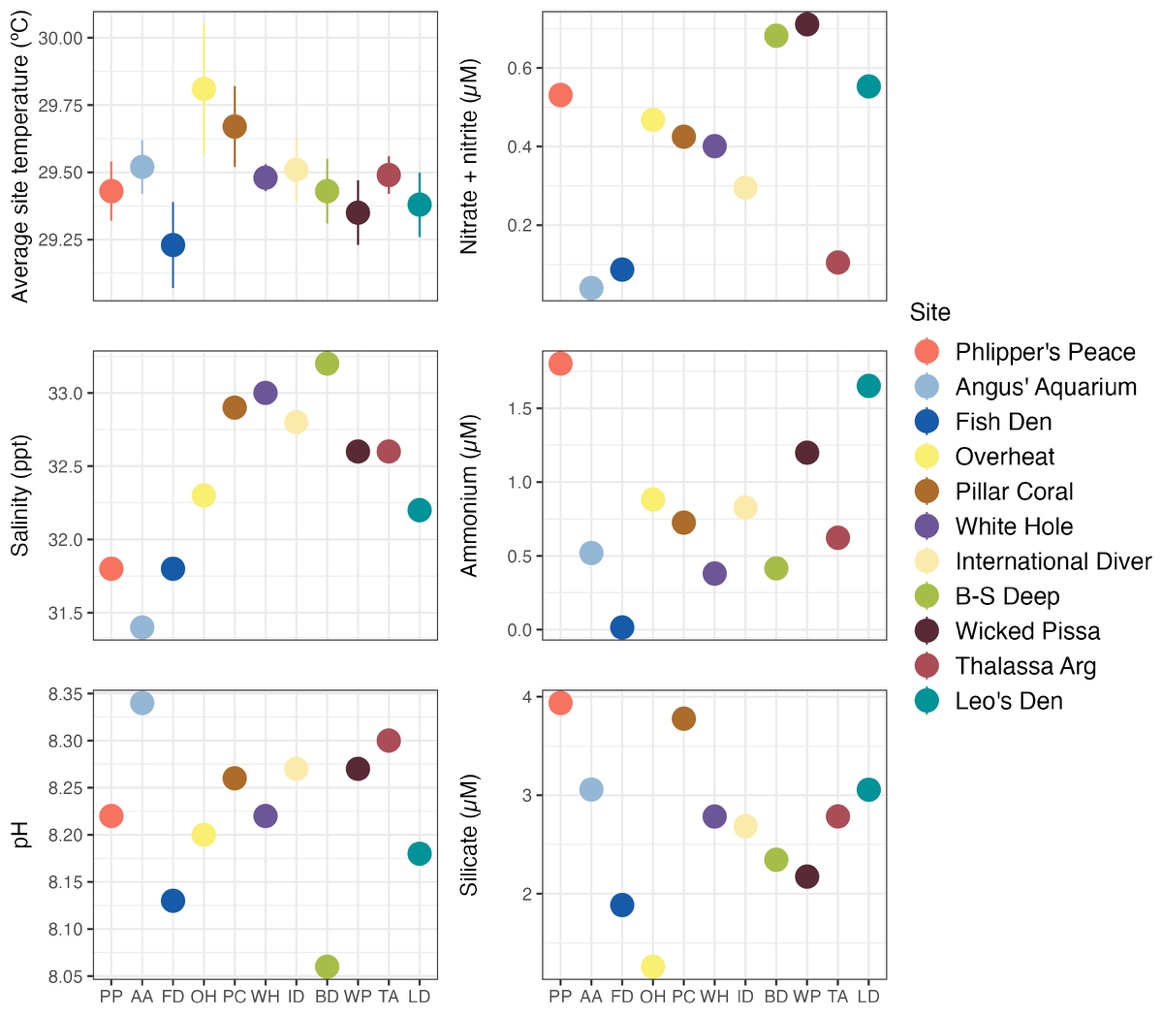

**Supplemental Figure S2.** Environmental characteristics at each site. In the average site temperature plot, points represent means and error bars represent ± standard deviation. In all other plots, points represent a single measured value. Sites are ordered by geography from west to east.

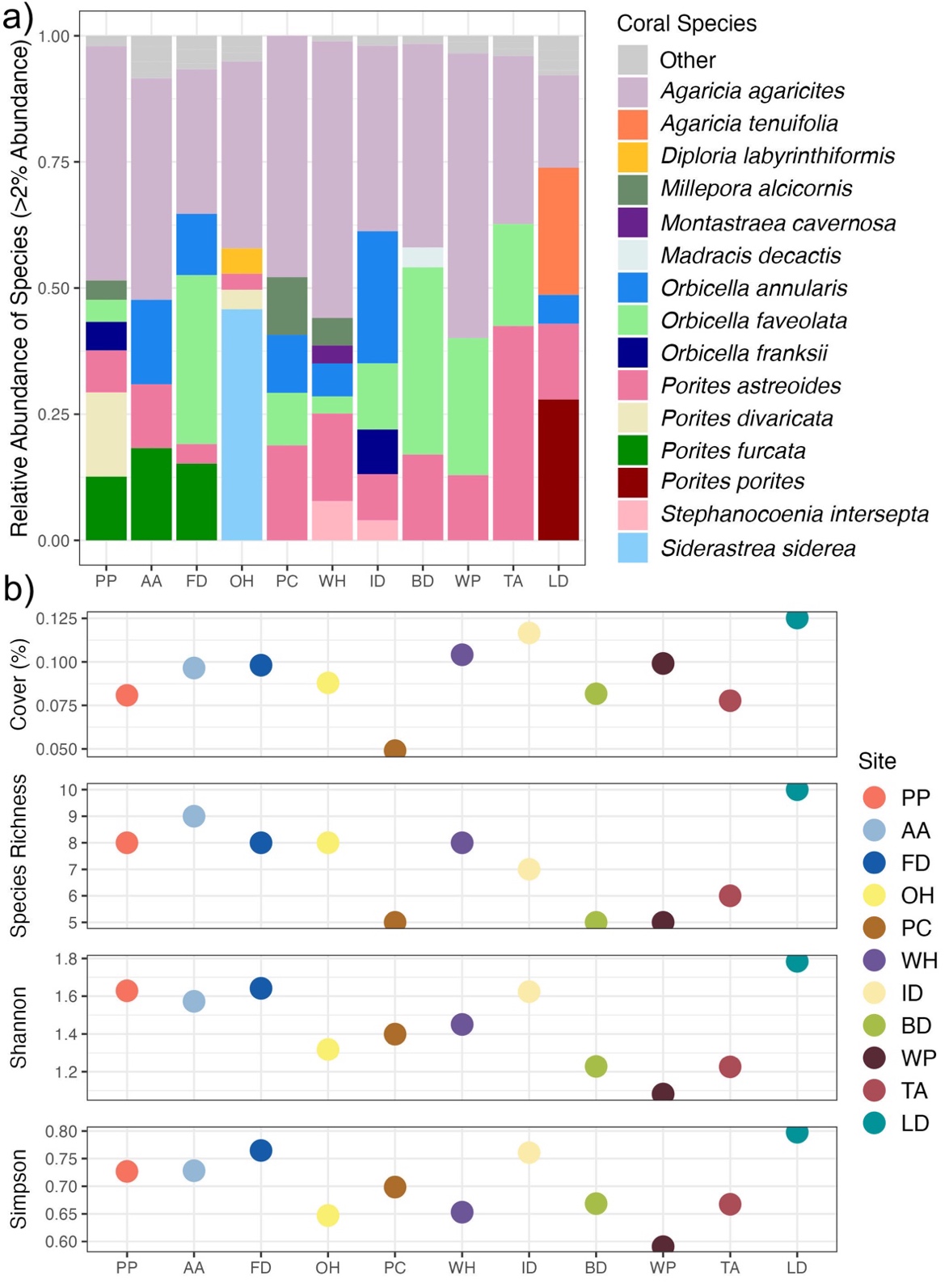

**Supplemental Fig S3.** Overview of coral a) community composition and b) alpha diversity. The bar plot demonstrates the relative abundance of coral species at each site. Each bar represents one site, and the “Other” category represents coral species present at <3% abundance at each site. The dot plots show coral percent cover, coral species richness, and the Shannon and Simpson diversity indices associated with coral communities at each site. Simpson diversity is presented as 1–D.

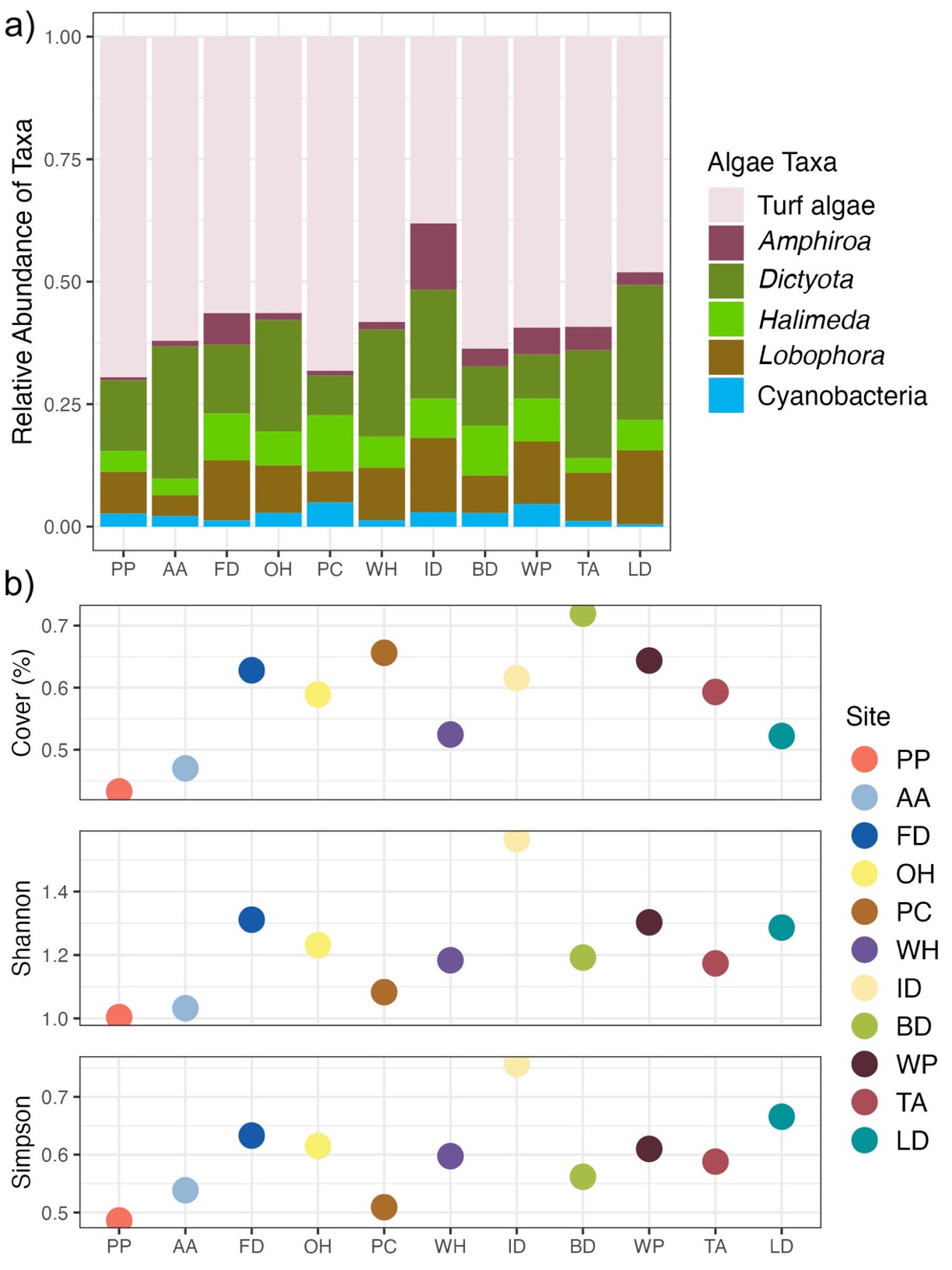

**Supplemental Figure S4.** Overview of macroalgae a) community composition and b) alpha diversity. The bar plot demonstrates the relative abundance of algal taxa at each site. Each bar represents one site. The dot plots show macroalgae percent cover and the Shannon and Simpson diversity indices associated with algae communities at each site. Simpson diversity is presented as 1–D.

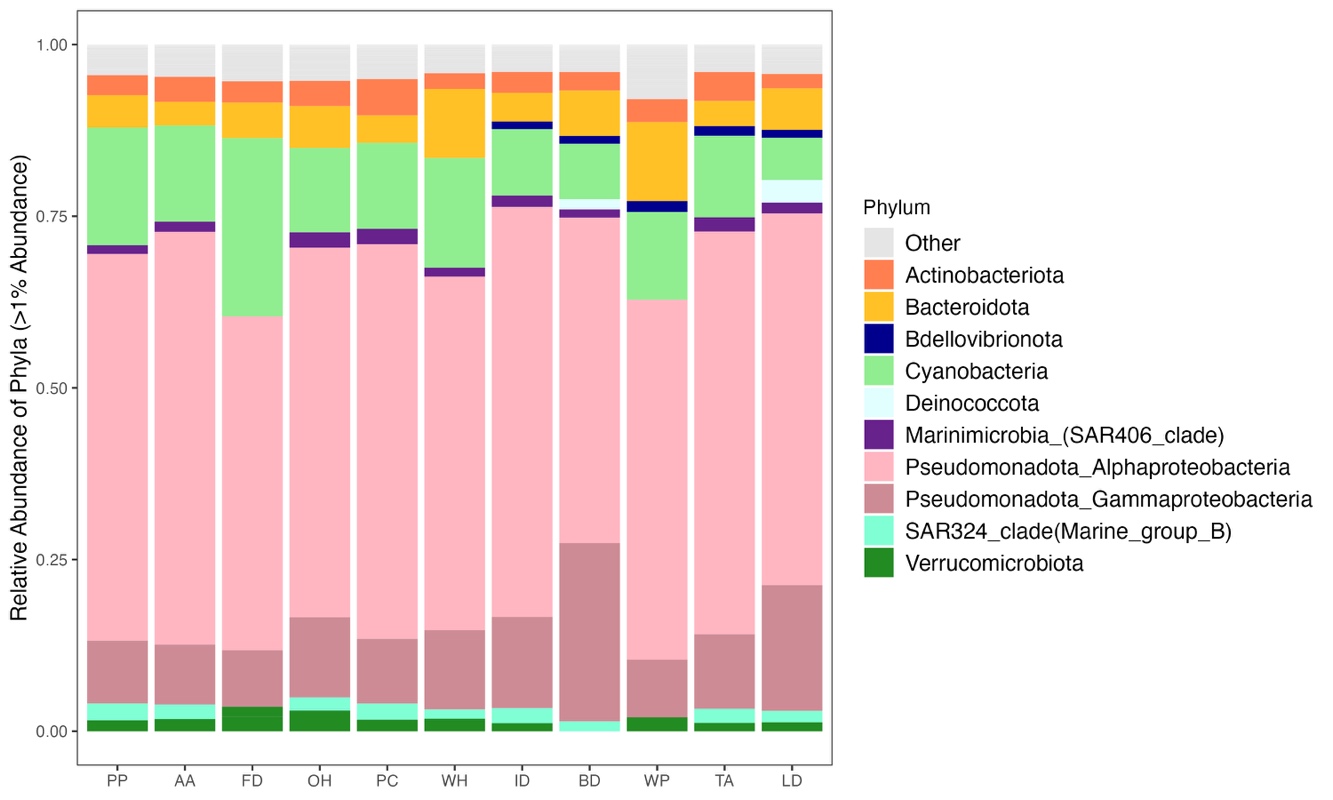

**Supplemental Figure S5.** Relative abundance of bacterial taxa in bacterioplankton samples. Each bar represents one reef site, colored by bacterial phylum, and the samples are ordered by geography. The “Other” group represents all taxa present at <1% abundance in each sample.

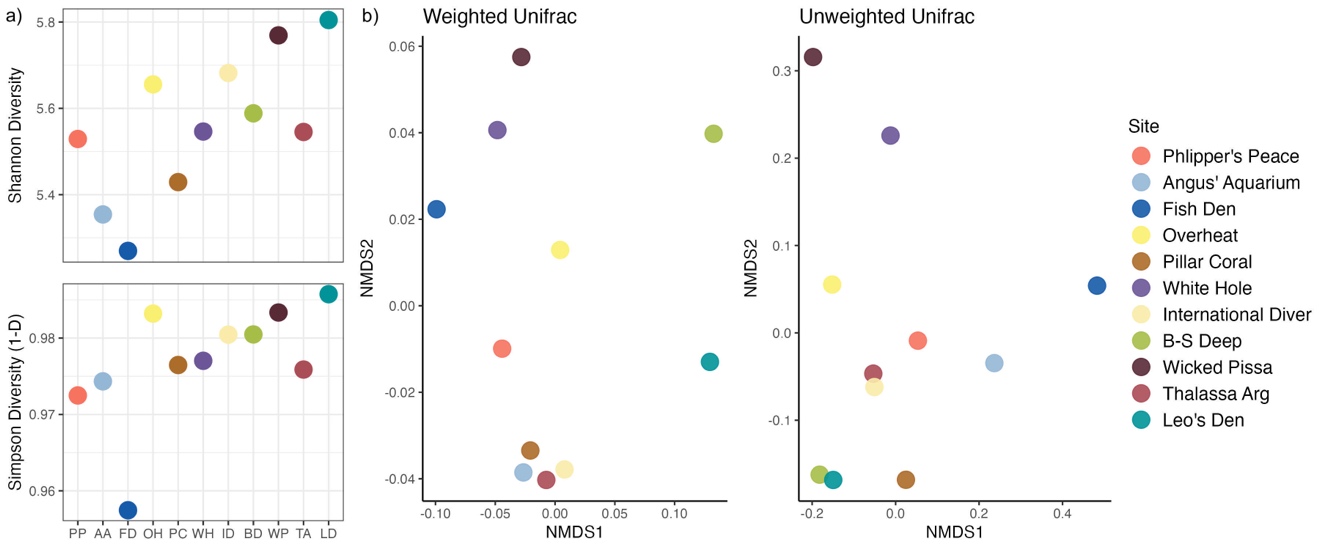

**Supplemental Figure S6.** a) Alpha and b) beta diversity of bacterioplankton communities. Simpson diversity is presented as 1–D, and points are ordered by geography from west to east in alpha diversity plots. For NMDS plots, distances were calculated with weighted and unweighted UniFrac metrics. Sites are ordered by geography from west to east.

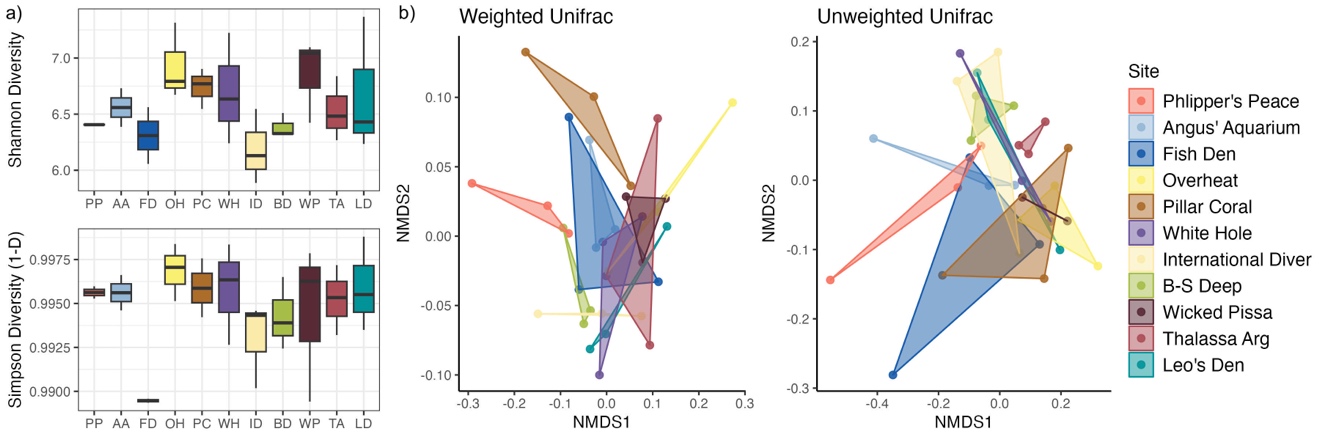

**Supplemental Figure S7.** a) Alpha diversity and b) community composition of CCA-associated biofilm communities by site. Simpson diversity is presented as 1–D, and boxplots are ordered by geography from west to east in alpha diversity plots. For NMDS plots, distances were calculated based on weighted and unweighted UniFrac metrics. Sites are ordered by geography from west to east.

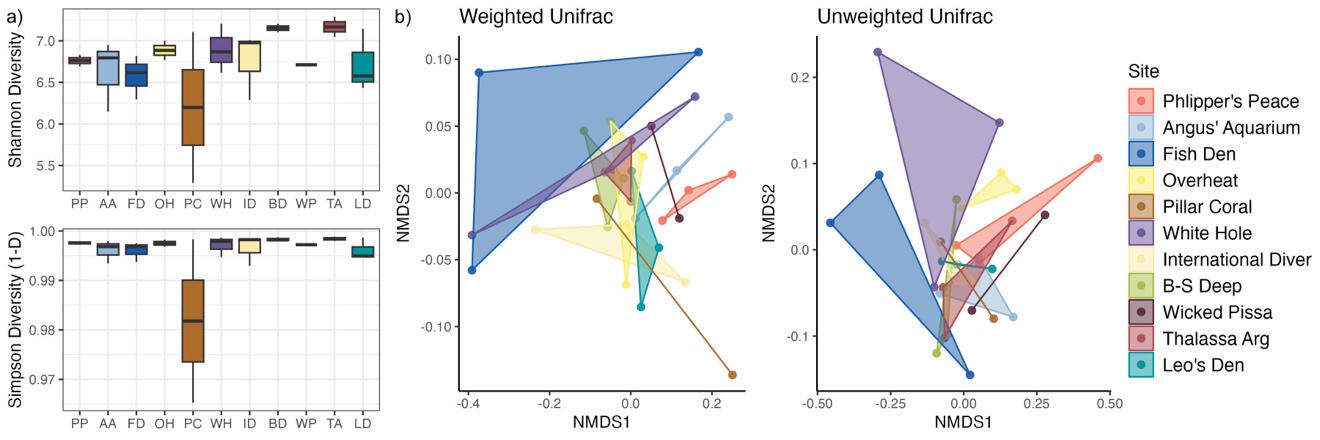

**Supplemental Figure S8.** a) Alpha diversity and b) community composition of carbonate-associated biofilm communities by site. Simpson diversity is presented as 1–D, and boxplots are ordered by geography in alpha diversity plots. For NMDS plots, distances were calculated based on weighted and unweighted UniFrac metrics. Sites are ordered by geography from west to east.
