## Supplemental Text for "Convergent stochastic assembly governs reef biofilm microbiomes across ecologically distinct benthic substrates"

Text S1:

CCA individuals were selected for sampling based on morphological similarities to minimize taxonomic variation and thus represent a single morphospecies. To sample benthic biofilm communities, a 10-oz claw hammer and sharpened stainless-steel tube were used to punch 1.6 cm-diameter cores from the reef substrate. The entire sample, including the stainless-steel tube and the core, was placed into a sterile collection bag (Whirl-Pak, Pleasant Prairie, WI) and transported to the RIMS laboratory on ice for immediate processing. In the RIMS dry laboratory, cores were carefully removed from tubes by gently tapping the tube on a sterilized countertop. The top surface of each core was swabbed using a high-retention microbial swab (Puritan Medical Products, Guilford, ME), and swabs were stored in DNA/RNA Shield (Zymo Research, Irvine, CA).

Additionally, at each site, a 1-L seawater sample was collected at 9-m depth for nutrient and microbial community analysis. In the RIMS laboratory, ~40 mL of each water sample was poured into labeled sterile 50-mL conical tubes and immediately frozen at –20ºC for nutrient analysis. The remaining volume from the water samples were syringe-filtered through 0.22-µm Sterivex filters (MilliporeSigma, Rockville, MD) to capture the bacterioplankton communities. The filters were removed from each filter unit using sterile channel lock pliers and a scalpel and stored in DNA/RNA Shield. All sample tubes with DNA/RNA Shield, including swabs and filters, were immediately stored at –20 ºC in the RIMS laboratory freezer. All samples were transported from RIMS to George Mason University in a cooler with blue ice, where filters and swabs were stored at –20 ºC and nutrient samples were stored at –80 ºC until further processing.

Photographs were taken for three replicate phototransects at each site to collect site-level coral and algae cover and composition data. Ten-meter transects were placed starting at 9 ± 1.5 m depth using a leaded line following the topography of the reef. Photographs were taken of a 0.25-m x 0.25-m quadrat every meter on alternating sides of the transect line, for a total of 10 quadrat photos per transect.

### Text S2:

Water samples for nutrient analysis were shipped to the Woods Hole Oceanographic Institute’s Nutrient Analytical Facility for quantification of ammonium, combined nitrate and nitrite, silicate, and orthophosphate using a SEAL Analytical AutoAnalyzer 3 HR (WHOI 2025). Orthophosphate concentrations were below the detection limit (<0.009 µM) at seven of 11 sites, so this nutrient was removed from further analysis. Phototransects were analyzed using Coral Point Count software to calculate percent cover and diversity metrics of stony coral and macroalgae (Kohler and Gill 2006). Benthic colonizers under 30 randomly distributed points on each photoquadrat were visually identified to the lowest possible taxonomic level using Humann and Deloach (2013). Coral and macroalgae cover and diversity metrics, including richness and Shannon and Simpson diversity indices, were based on all 30 quadrat photos from each site to calculate a composite site-level value.

### Text S3:

Genomic DNA was extracted from the swab and filter samples using QIAGEN DNeasy PowerBiofilm Kits (Qiagen, Germantown, MD) following the manufacturer’s protocol with the following modifications. For each sample, the entire swab or filter was transferred into a PowerBiofilm Bead Tube using ethanol- and flame-sterilized forceps. Microbial materials remaining in the DNA/RNA Shield were pelleted by centrifugation, the DNA/RNA Shield was poured off, and the pellet was resuspended and added to the PowerBiofilm Bead Tube according to the protocol. Bead beating occurred on an Omni Bead Ruptor 24 for 30 s at 1.6 m/s speed for both sample types. DNA was eluted in 100 µL of diethylpyrocarbonate (DEPC)-treated water (Ambion, Austin, TX), and extracted DNA was stored at –20 ºC. Following extraction, genomic DNA was quantified using a Qubit 2.0 fluorometer and Qubit dsDNA High Sensitivity Assay Kit (Invitrogen, Waltham, MA) following the manufacturer’s protocol.

### Text S4:

FASTQ files provided by IMR were processed with the package DADA2 (version 1.26.0) in RStudio (R version 4.2.1) to remove primers, length-filter reads to those between 1000–1600 nt, and quality-filter reads to remove sequences with ambiguous bases or greater than 20 expected errors (Callahan et al. 2016). Filtered sequences were then imported into the QIIME2 bioinformatic pipeline (version 2024.10, (Bolyen et al. 2019, QIIME2 Development Team 2023). Briefly, sequences were dereplicated, clustered into operational taxonomic units (OTUs) with a 99% identity cut-off, and chimeric sequences were identified and removed. Taxonomy was assigned using the SILVA 138 reference database using the classify-sklearn option with a 90% confidence cut-off (Quast et al. 2013, Bokulich et al. 2018). OTUs were then aligned to representative sequences using MAFFT (Katoh 2002) and used to create a phylogenetic tree with FastTree (Price et al. 2010).

### Text S5:

To investigate the relative influences of selection, dispersal, and drift on the biofilm communities, an analysis was performed with the program Infer Community Assembly Mechanisms by Phylogenetic bin-based null model analysis (“iCAMP analysis”), developed by Ning et al. (2020) based on a previous statistical framework (Stegen et al. 2013), implemented using the iCAMP package (version 1.5.12). OTUs were grouped by genus and then binned based on phylogenetic relatedness. First, abundance-weighted 𝛽-mean-nearest taxon distance (𝛽MNTD) was calculated for each pair of bins and compared to a null model based on 999 randomly assembled communities to determine the 𝛽-nearest taxon index (𝛽NTI). Values of |𝛽NTI| > 1.96 indicate significantly more or less phylogenetic turnover than expected by the null model and were attributed to selection (Stegen et al. 2012, 2013), with positive values assigned to heterogeneous selection and negative values assigned to homogeneous selection (Stegen et al. 2015). For pairwise comparisons with |𝛽NTI| ≤ 1.96, an index based on Bray-Curtis-based Raup-Crick distances (RCBray) was calculated for each bin pair and compared to a null model based on 999 randomly assembled communities. When |RCBray| > 0.95, there was significantly more or less OTU turnover than expected by drift alone (i.e., null model), which was attributed to dispersal (Stegen et al. 2013), with positive values assigned to dispersal limitation and negative values assigned to homogenizing dispersal (Stegen et al. 2015). The remaining bin pairs with |𝛽NTI| ≤ 1.96 and |RCBray| ≤ 0.95 were assigned to ecological drift, but it should be noted that this category may also include influence from stochastic diversification, weak selection, and weak dispersal (Ning et al. 2020).

### Text S6:

Environmental parameters were variable among sites, with no clear spatial trends (Supp. Fig. S2). Additionally, differences in environmental conditions were not significantly correlated with geographic distance (Mantel test, p = 0.39). The coral communities at most sites were dominated by *Agaricia agaricites* (40.4% ± 11.3%, mean ± standard deviation), but community composition was variable among sites (Supp. Fig. S3a). Coral cover ranged from 4.9–12.5% among sites, and alpha diversity of coral communities was variable among sites (Supp. Fig. S3b). Coral cover and alpha diversity were both highest at LD. Coral cover was lowest at PC, but diversity was lowest at WP. Algal communities at all sites were dominated by turf algae (57.9% ± 9.0%, Supp. Fig. S4a). Algae cover was high, ranging from 43.3% at PP to 71.9% at BD, but algae diversity was variable, highest at ID and lowest at PP (Supp. Fig. S4b). There was a weak but significant correlation between coral community distance and geographic distance, suggesting that sites that are closer together have more similar coral communities (Mantel test, r = 0.23, p = 0.04), but there was no significant correlation between algal composition and geographic distance (Mantel test, p = 0.15).

Bacterioplankton communities were analyzed at all sites to assess the influence of bacterioplankton community composition on biofilm community composition. Long-read amplicon sequencing of the 16S rRNA gene of 11 water samples resulted in 645,914 raw reads, with 394,474 reads remaining after quality filtering, corresponding to 38,833 OTUs. OTUs that were identified as mitochondrial (n = 1,815) or chloroplast (n = 2,182) DNA were removed, as well as four OTUs found in the negative control, leaving 367,589 reads and 34,832 ASVs in the final dataset used for analysis.

Bacterioplankton community composition at the phylum level was very consistent among sites and was dominated by Alphaproteobacteria (Pseudomonadota), but other phyla present in every sample at >1% abundance include Actinobacteriota, Bacteroidota, Cyanobacteria, and Gammaproteobacteria (Pseudomonadota, Supp. Fig. S5). Bacterioplankton communities were highly diverse, with some variation in Shannon and Simpson diversity among sites, and community diversity increased geographically from west to east (Supp. Fig. S6a). Community composition was variable among sites at the OTU level. Although some sites were more similar than others, there were no distinct geographic patterns according to weighted or unweighted UniFrac distances (Supp. Fig. S6b). This is further supported by non-significant correlations between genetic and geographic distances in bacterioplankton communities according to both weighted (Mantel test, p = 0.40) and unweighted (p = 0.20) UniFrac distances. There was also no significant correlation between bacterioplankton community composition and environmental conditions according to weighted (Mantel test, p = 0.57) or unweighted (p = 0.61) UniFrac distances.

### Text S7:

In CCA-associated biofilm communities, there were no significant differences in Shannon or Simpson diversity (ANOVA, p > 0.19 for both, Supp. Fig. S7a) or beta dispersion based on weighted and unweighted UniFrac distances (PERMDISP, p > 0.84 for both) among sites. There were significant differences in community composition among sites based on weighted and unweighted UniFrac distances (PERMANOVA, p < 0.04 for both, Supp. Fig. S7b), but none of the pairwise comparisons were significant (p > 0.1 for all). In carbonate-associated biofilm communities, there were no significant differences in Shannon or Simpson diversity (ANOVA, p > 0.37 for both, Supp. Fig. S8a), beta dispersion (weighted and unweighted UniFrac, PERMDISP, p > 0.41 for both), or community composition (weighted and unweighted UniFrac, PERMANOVA, p > 0.11 for both, Supp. Fig. S8b) among sites. Additionally, neither CCA- nor carbonate-associated biofilm community composition were significantly correlated with geographic distance between samples based on weighted or unweighted UniFrac distances (Mantel tests, p > 0.22 for all).

Bolyen, E., J. R. Rideout, M. R. Dillon, N. A. Bokulich, C. C. Abnet, G. A. Al-Ghalith, H. Alexander, E. J. Alm, M. Arumugam, F. Asnicar, Y. Bai, J. E. Bisanz, K. Bittinger, A. Brejnrod, C. J. Brislawn, C. T. Brown, B. J. Callahan, A. M. Caraballo-Rodríguez, J. Chase, E. K. Cope, R. Da Silva, C. Diener, P. C. Dorrestein, G. M. Douglas, D. M. Durall, C. Duvallet, C. F. Edwardson, M. Ernst, M. Estaki, J. Fouquier, J. M. Gauglitz, S. M. Gibbons, D. L. Gibson, A. Gonzalez, K. Gorlick, J. Guo, B. Hillmann, S. Holmes, H. Holste, C. Huttenhower, G. A. Huttley, S. Janssen, A. K. Jarmusch, L. Jiang, B. D. Kaehler, K. B. Kang, C. R. Keefe, P. Keim, S. T. Kelley, D. Knights, I. Koester, T. Kosciolek, J. Kreps, M. G. I. Langille, J. Lee, R. Ley, Y.-X. Liu, E. Loftfield, C. Lozupone, M. Maher, C. Marotz, B. D. Martin, D. McDonald, L. J. McIver, A. V. Melnik, J. L. Metcalf, S. C. Morgan, J. T. Morton, A. T. Naimey, J. A. Navas-Molina, L. F. Nothias, S. B. Orchanian, T. Pearson, S. L. Peoples, D. Petras, M. L. Preuss, E. Pruesse, L. B. Rasmussen, A. Rivers, M. S. Robeson, P. Rosenthal, N. Segata, M. Shaffer, A. Shiffer, R. Sinha, S. J. Song, J. R. Spear, A. D. Swafford, L. R. Thompson, P. J. Torres, P. Trinh, A. Tripathi, P. J. Turnbaugh, S. Ul-Hasan, J. J. J. van der Hooft, F. Vargas, Y. Vázquez-Baeza, E. Vogtmann, M. von Hippel, W. Walters, Y. Wan, M. Wang, J. Warren, K. C. Weber, C. H. D. Williamson, A. D. Willis, Z. Z. Xu, J. R. Zaneveld, Y. Zhang, Q. Zhu, R. Knight, and J. G. Caporaso. 2019. Reproducible, interactive, scalable and extensible microbiome data science using QIIME 2. Nature Biotechnology 37:852–857.

Callahan, B. J., P. J. McMurdie, M. J. Rosen, A. W. Han, A. J. A. Johnson, and S. P. Holmes. 2016. DADA2: High-resolution sample inference from Illumina amplicon data. Nature Methods 13:581–583.

Humann, P., and N. Deloach. 2013. Reef coral identification: Florida, Caribbean, Bahamas. Third edition. New World Publications, Jacksonville, FL.

Katoh, K. 2002. MAFFT: A novel method for rapid multiple sequence alignment based on fast Fourier transform. Nucleic Acids Research 30:3059–3066.

Kohler, K. E., and S. M. Gill. 2006. Coral Point Count with Excel extensions (CPCe): A Visual Basic program for the determination of coral and substrate coverage using random point count methodology. Computers & Geosciences 32:1259–1269.

Ning, D., M. Yuan, L. Wu, Y. Zhang, X. Guo, X. Zhou, Y. Yang, A. P. Arkin, M. K. Firestone, and J. Zhou. 2020. A quantitative framework reveals ecological drivers of grassland microbial community assembly in response to warming. Nature Communications 11:4717.

Price, M. N., P. S. Dehal, and A. P. Arkin. 2010. FastTree 2 – Approximately maximum-likelihood trees for large alignments. PLoS ONE 5:e9490.

QIIME2 Development Team. 2023. Overview of QIIME2 plugin workflows. https://docs.qiime2.org/2023.9/tutorials/overview/#let-s-get-oriented-flowcharts.

Quast, C., E. Pruesse, P. Yilmaz, J. Gerken, T. Schweer, P. Yarza, J. Peplies, and F. O. Glöckner. 2013. The SILVA ribosomal RNA gene database project: Improved data processing and web-based tools. Nucleic Acids Research 41:D590–D596.

Stegen, J. C., X. Lin, J. K. Fredrickson, X. Chen, D. W. Kennedy, C. J. Murray, M. L. Rockhold, and A. Konopka. 2013. Quantifying community assembly processes and identifying features that impose them. The ISME Journal 7:2069–2079.

Stegen, J. C., X. Lin, J. K. Fredrickson, and A. E. Konopka. 2015. Estimating and mapping ecological processes influencing microbial community assembly. Frontiers in Microbiology 6:370.

Stegen, J. C., X. Lin, A. E. Konopka, and J. K. Fredrickson. 2012. Stochastic and deterministic assembly processes in subsurface microbial communities. The ISME Journal 6:1653–1664.

WHOI. 2025. Current Rates. https://web.whoi.edu/nutrient/current-rates/.
